# Pleiotropy alleviates the fitness costs associated with resource allocation trade-offs in immune signaling networks

**DOI:** 10.1101/2023.10.06.561276

**Authors:** Reese Martin, Ann.T. Tate

## Abstract

Many genes and signaling pathways within plant and animal taxa drive the expression of multiple organismal traits. This form of genetic pleiotropy instigates trade-offs among life-history traits if a mutation in the pleiotropic gene improves the fitness contribution of one trait at the expense of another. Whether or not pleiotropy gives rise to conflict among traits, however, likely depends on the resource costs and timing of trait deployment during organismal development. To investigate factors that could influence the evolutionary maintenance of pleiotropy in gene networks, we developed an agent-based model of co-evolution between parasites and hosts. Hosts comprise signaling networks that must faithfully complete a developmental program while also defending against parasites, and trait signaling networks could be independent or share a pleiotropic component as they evolved to improve host fitness. We found that hosts with independent developmental and immune networks were significantly more fit than hosts with pleiotropic networks when traits were deployed asynchronously during development. When host genotypes directly competed against each other, however, pleiotropic hosts were victorious regardless of trait synchrony because the pleiotropic networks were more robust to parasite manipulation, potentially explaining the abundance of pleiotropy in immune systems despite its contribution to life history trade-offs.

## Introduction

Life-history trade-offs arise when two or more traits cannot simultaneously increase their contribution to organismal fitness. Trade-offs can occur between discrete traits, such as organ systems competing over resources during development (Nijhout and Emlen 1998) or within traits, as with the careful balance between autoimmunity and defense against pathogens achieved by the immune system (Graham et al. 2022). Life-history trade-offs are evolutionarily interesting because they highlight traits that experience conflicting selective pressures (Sinervo and Svensson 1998), leading to the maintenance of morphological diversity within species or even driving speciation events, as when conflict in body size drives the increasing reproductive isolation between ‘normal’ and ‘dwarf’ arctic charr (Parker et al. 2001).

Resource allocation trade-offs are a special case of life-history trade-offs that occur when a limited resource must be distributed among multiple traits, ultimately resulting in at least one trait receiving a less than optimal allotment. Often the resource in demand is some form of energetic currency that must be divvied up between intensive traits like reproduction, growth (Bleu et al. 2012), or immunity (Husak et al. 2017). The allocation of energy to fuel an immune system can have lasting consequences for fitness-associated traits, as when thale cress (*Arabidopsis thaliana*) sacrifices fecundity to defend against the bacterium *Pseudomonas syringae* (Tian et al. 2003). Allocation issues can also impose more transient consequences, as when intense bouts of energy expenditure leave hosts periodically susceptible to infection and severe disease (Nieman and Pedersen 1999; Adamo and Parsons 2006). Resource allocation trade-offs could even affect network robustness, the fundamental ability of a signaling network to consistently produce a response in the face of manipulation or interference, particularly when robust networks require more energetic (Bullmore and Sporns 2012) and component (Kitano 2004) resources than non-robust networks.

Timing plays a critical role in resource allocation trade-offs (Nieman and Pedersen 1999; Adamo and Parsons 2006), with many trade-offs only becoming evident when two (or more) traits are engaged at the same time. A classic example of this phenomenon manifests in female insects that try to reproduce after infection or as they try to fend off an infection following mating. The former leads to fewer viable offspring and the later leading to decreased infection survival (Schwenke et al. 2016). While this trade-off between immunity and fecundity is a consequence of a global metabolic reorganization, trade-offs can also stem from limited amounts of a single product. Such a trade-off is seen with ribosome availability limiting cell growth, especially when cells are required to synthesize proteins that are not growth-related (Carrera et al. 2011). Thus, the synchrony of deploying two traits in space and time, as in specific tissues or developmental stages, is likely to modulate the magnitude and slope of resource allocation trade-offs.

Pleiotropy, the phenomenon where a gene or gene product mediates the expression of two or more traits, can give rise to trade-offs through genetic and evolutionary constraints, resource allocation conflicts, or potentially both at the same time, depending on the synchrony of trait expression. Pleiotropy is expected to evolve as organisms use extant genetic material to develop novel traits, as has been observed theoretically (Lenski et al. 2003) and empirically through the genomic study of trait architecture (Armbruster et al. 2009). Pleiotropy may also arise as a result of selection on co-varying traits (Cheverud 1996), but pleiotropy is not always beneficial, especially across environments and signaling conditions (Kinsler et al. 2020). Antagonistic pleiotropy arises when the traits a pleiotropic gene participates in cannot simultaneously increase their contributions to organismal fitness, such as the trade-off between larval survival and adult size observed in the fruit fly *Drosophila melanogaster* (Bochdanovits and de Jong 2004). Such antagonistic pleiotropy is a phenomenon found across the tree of life, with examples found in plants (Scarcelli et al. 2007; Sadhukhan et al. 2021), animals (Bochdanovits and de Jong 2004), prokaryotes (Reyes et al. 2013; Moleres et al. 2018), and even viruses (Bedhomme et al. 2015). Antagonistic pleiotropy can arise due to a trade-off in function; for example, cardiac insulin like growth factor is vital for juvenile cardiac development in mice, *Mus musculus*, but leads to age-associated cardiac degeneration (Abdellatif et al. 2023).

Antagonistic pleiotropy can also arise as a matter of resource allocation, where the resource in need of allocation could be the pleiotropic protein or signaling pathway module, a substrate or output, or the energetic currency necessary to fuel the pleiotropic traits (Hughes and Leips 2017). Pleiotropic genes are abundant in the immune system (Sivakumaran et al. 2011) and pleiotropic immune genes have been shown to evolve more slowly than non-pleiotropic immune genes (Williams et al. 2023). An example of antagonistic pleiotropy in immunity manifests in the reduced melanization potential of Spongy moth (*Lymantria dispar*) larvae immediately following molting (McNeil et al. 2010). In this system melanin is used for cuticle tanning as well as encapsulating pathogens, but its production is limited because of the potential for self-inflicted damage from reactive intermediaries and cytotoxic quinines (González-Santoyo and Córdoba-Aguilar 2012), leading to a trade-off when both tanning and immune defense are attempted simultaneously. This example presents compelling evidence that changes in the timing of signaling network activity may affect the severity of trade-offs between traits, with trade-offs becoming apparent when signaling is synchronous, and disappearing when signaling activity is asynchronous. Therefore, we chose to investigate the interaction of resource availability, effector module pleiotropy, and signaling synchrony over organismal development.

The melanization trade-off in *L*.*dyspar* is just one example of how the use of one pathway, module, or effector for two simultaneous purposes is likely to generate an outsized allocation or optimization trade-off among pleiotropic traits. Table 1 contains several more examples of pleiotropy in signaling networks; some of these present clear allocation problems, and some seem largely benign, but still pose the question: Does trait synchrony matter for the evolution and maintenance of pleiotropy? Does resource allocation alter this answer? We hypothesize that pleiotropic effectors are most detrimental to organismal fitness when signaling is synchronous because the simultaneous expression of both traits will exacerbate resource conflict, particularly when resources are scarce.

**Table 1:** Biological examples of pleiotropic resource allocation trade-offs.

| Gene/pathway | Timing of trade-off | Traits | System | parasite | Summary | References |
| --- | --- | --- | --- | --- | --- | --- |
| InR | Asynch. | *Growth/fecundity and lifespan/stress tolerance* | *D. melanogaster*<br>*Common fruit fly* | N/A | The Insulin-like Receptor (InR) in *D. melanogaster* is associated with trade-offs between fecundity and development time vs lifespan and stress tolerance. | (Paaby et al. 2014) |
| RXFP2 | Asynch. | *Fecundity and organismal survival* | *Ovis aries*<br>*Domestic sheep* | N/A | Relaxin-like receptor 2 (RXFP2) is associated with a trade-off between reproductive success and year over year survival. | (Johnston et al. 2013) |
| RPM1 | Asynch. | *Infection resistance and seed production* | *A. thaliana*<br>*Thale cress* | *P. syringae* | The *RPM1* gene confers resistance to *Pseudomonas syringae* in *A. thaliana* but reduces reproductive fitness. | (Bisgrove et al. 1994; Tian et al. 2003) |
| TNFα | Asynch. | *Immune organ development and offspring quality quantity ratio* | TNF-KO Mouse<br>*Mus musculus* | N/A | TNFα is critical for immune homeostasis and maintains the quantity vs. quality trade-off in mouse litters. | (Maslennikova et al. 2019) |
| (HIF)-1α | Synch | *Immune defense and wound healing* | (HIF)-1α KO mouse<br>*Mus musculus* | Group A *streptococcus* strain 5448 | Hypoxia-inducible transcription factor (HIF)-1α allows NK cells to mediate a metabolic trade-off between wound healing behavior and bacterial defense. | (Sobecki et al. 2021) |
| Juvenile Hormone (JH) and 20 hydroxy ecdysone (20E) | Synch. | *AMP production and egg production* | Review of findings across insects | N/A | JH is an antagonist of immune function while 20E is a potentiator of immunity. The balance between JH and 20E is vital for egg maturation. | (Schwenke et al. 2016) |
| ProPO | Synch. | *Pathogen melanization and cuticle tanning* | *Lymantria dispar*<br>*Spongy moth* | *LDMNPV* | Phenoloxidase is used to both tan insect cuticles and melanize pathogens, this leads to lowered resistance to infection immediately following molting. | (McNeil et al. 2010) |
| Toll | Synch. | *AMP production and growth* | Gal4/UAS *D. melanogaster* constitutively expressing Toll<sup>10b</sup><br>*Common fruit fly* | N/A | Suppression of insulin signaling following the activation of Toll signaling in the fat body of *Drosophila melanogaster* suppresses allocation of resources to growth. | (DiAngelo et al. 2009) |

To address this hypothesis, we developed a theoretical model of signaling network co-evolution featuring host and parasite populations. Drawing inspiration from the partially pleiotropic Toll and ProPO pathways in insects (Table 1), hosts are defined by a pair of signaling networks, one developmental and the other immune. Signaling networks that control each trait can be fully independent (independent effector hosts) or feature two partially decoupled signaling networks that share a single effector or downstream module (shared effector hosts). Host life-history includes an immature stage where they are subjected to developmental signals and an adult stage where there is no developmental signaling.

Parasites are defined by a single network that allows them to interfere with host immune signaling, an common form of host-parasite interaction that is thought drive host immune signaling network evolution (Schmid-Hempel 2008). In our model, hosts defend against parasitic infection during their developmental period or following the end of development, creating a synchronous or asynchronous signaling environment. Hosts evolve in one of three resource conditions (scarce, plentiful, alternating) which limit the actions of proteins in their signaling networks. We use this modeling framework to investigate the conditions that favor the evolutionary maintenance of pleiotropy despite the proximate and ultimate constraints it imposes on adaptation, revealing benefits to pleiotropy that only manifest in heterogeneous populations.

## Methods

We adapted a framework originally designed to model the evolution of generic signaling networks (Soyer et al. 2006) and subsequently modified to study immune signaling network robustness (Salathé and Soyer 2008), the evolvability of immune responses during co-evolution (Schrom et al. 2018), and the interaction between pleiotropy and immune response dynamics during co-evolution (Martin and Tate 2023). Here we have revised the model to include two signaling networks that can function independently but have a shared resource pool. While these modifications were originally made to reflect pleiotropy between immune and developmental processes, the model itself is generalizable to any set of pleiotropic traits. Aside from sharing effector modules (downstream pleiotropy), we also allowed hosts to develop internetwork connections (upstream pleiotropy) to study the fitness effects of pleiotropic interactions between two networks as well as the conditions that promoted the creation and maintenance of such connections. For this work we elected to use a simulation of network evolution rather than analytically determining optimum traits because it allows us to evaluate the ‘evolvability’ of an evolutionary endpoint as well as avoid potential biases in results from our assumptions about what traits or trait combinations are optimal.

For the purposes of this project, we broadly defined an effector as a module of genes (e.g. the intracellular portion of the Toll pathway), a regulatory element, or output (e.g. melanin from phenoloxidase). Hosts with a shared (pleiotropic) effector had a single effector controlled by both signaling networks and hosts with independent effectors had two independent effectors each controlled by a distinct signaling network. In all simulations, each host was initially composed of two networks, with independent effector hosts having a detector, three signaling proteins, and an effector in each network, and shared effector hosts having a detector and three signaling proteins in each network, but a single effector shared by both networks.

Connections between proteins and the regulatory behavior of those connections were randomized during the initiation of a simulation. Host creation parameters were derived from previous work concerning agent based models of signaling network evolution (Soyer et al. 2006; Salathé and Soyer 2008; Schrom et al. 2018; Martin and Tate 2023). Host mutations occurred during reproduction and resulted in the duplication or deletion of signaling proteins, or the addition, deletion, or modification of regulatory interactions between proteins in the network. The only restrictions to this evolutionary process were that detectors and effectors were not allowed to develop a direct connection and could not be deleted. Hosts were required to maintain at least one signaling protein, preventing them from evolving an entirely non-functional immune network. Host fitness was expressed as a function of immune effector abundance, cumulative parasite abundance, and how closely they matched a developmental input (see “fitness calculation” section, below). Parasite fitness was derived from cumulative parasite abundance, a proxy for transmission potential.

To evaluate the evolutionary implications of hosts using a shared effector rather than multiple independent effectors we designed two major classes of simulation: co-evolutionary, designed to establish a fitness landscape to identify resource and signaling conditions that favored independent or shared effector hosts, and competitive simulations, designed to evaluate the competitive fitness of shared and independent effector hosts against each other. During co-evolutionary simulations, a population of hosts (n = 500, either entirely shared effector hosts or entirely independent effector hosts) evolved for 1000 generations with a population of parasites (n = 250). The initial hosts and parasites in a population can be thought of as founder ‘genomes’, seeding the simulation with initial genetic variance for evolutionary processes to act on. Each host had a 50% chance of being infected by a parasite in each generation.

Competitive simulations consisted of two phases: independent evolution and competition. During the independent evolution phase of a competitive simulation a population of hosts with independent effectors (n = 250) and a population of hosts with shared effectors (n = 250) were allowed to co-evolve with parasite populations (n = 125) in isolation from the other host population. As in the independent evolution case, the initial hosts and parasites can be thought of as being a founding ‘genotype’ from which a successful lineage may descend. After 250, 500, or 1000 generations of independent evolution, the simulation entered the competition stage, combining the host populations along with their parasites.

The competition ended when one population died out entirely or 1000 generations had passed with no winner (draw). Host and parasite population size were chosen to balance initial population variability, a trait shown to affect evolutionary responses (Ørsted et al. 2019), against computational resources. The number of generations allowed for evolution was chosen to balance the degree of population fitness increases in later generations against the time necessary to complete a simulation. These simulations allowed us to further evaluate the costs and benefits associated with sharing an effector between multiple traits.

### Simulations of independent evolution

Evolutionary simulations were carried out in the Julia programming language (v 1.6) (Bezanson et al. 2017). Each simulation is composed of generations which can be broken down into the following discrete events. The initialization step happened only once per simulation.

#### 1. Initialization

a population of 500 shared effector or independent effector hosts is generated. All independent effector hosts start with a detector, three signaling proteins, and an effector for both their immune and their developmental signaling networks. All shared effector hosts start with a detector and three signaling proteins for their immune and developmental signaling networks but share a single effector. Connections are established between proteins in a signaling network at random, with each valid connection in the network having a 50% chance of occurring. Each connection is assigned a randomly selected regulatory value from the interval [-1,1]. A population of 250 parasites is also generated at the start of each simulation. Parasites are designated by a target (restricted to signaling proteins) and regulatory behavior on that target (restricted to the interval [-1,1]).

#### 2. Host equilibrium

Each protein P in an immune network has a total concentration equal to 1, with the active portion denoted P_i_^*^ and inactive portion denoted *P*_*i*_, ([P_i_^*^] + [*P*_*i*_] = 1). When determining the effects of protein P on other proteins in the network only the active portion is considered. During infection, changes in parasite abundance are calculated as though it was another protein in the network. The change in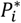 is determined by the regulatory action on *P*_*i*_ defined:

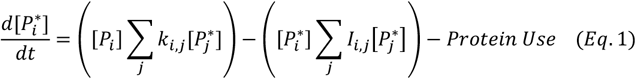

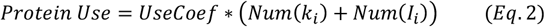

Where *k*_*i,j*_ are the upregulatory coefficients from protein *P*_*j*_ to protein *P*_*i*_, *I*_*i,j*_ are the downregulatory coefficients from protein *P*_*j*_ to protein *P*_*i*_. Here upregulatory connections are those that increase the abundance of the target protein and downregulatory connections are those that decrease target protein abundance. The term *Protein Use* describes the inactivation of some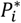 owing to the action of *P*_*i*_ on other proteins in the network, for all simulations *UseCoef* = .01 so for each interaction in a timestep a protein P_i_’s total active portion, [P_i_^*^], decreased by .01. A larger value would necessitate fewer interacting partners or a greater upregulation of protein P_i_ to maintain [P ^*^].

Starting from initial concentrations of P^*^ = .5 for all proteins in the network, P* for each protein at time step *t+1* is calculated numerically using Equation (1) for 5 time-steps to allow host signaling to reach equilibrium.

#### 3. Development

Following the equilibrium period, hosts begin receiving a developmental signal defined by the following equation 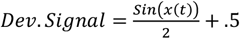, where *x*(t) are the sequential values in *range* (0,8π) at time-step t. Developmental signaling occurred for 50 time-steps in all simulations. This signaling dynamic was chosen to mimic a pulsed or periodic developmental signal used to coordinate periodic events like periodic segmentation, a process common in both vertebrates and invertebrates (Sonnen and Janda 2021).

#### 4. Infection

In each generation, 50% of hosts were selected at random for infection by a randomly selected parasite. Due to population size differences this means that all parasites infected a host in each generation of a simulation. Major results are presented with infection following development, so infection during asynchronous simulations occurred in a 50 timestep window following the end of the developmental period. For symmetry we also conducted a subset of simulations where developmental signaling took place following infection; refer to ‘Host Life’ below for details. In synchronous simulations, hosts were infected at a randomly selected point during the developmental period. Parasites are treated as an additional protein in the host immune network with an upregulatory connection of 1 to the host detector, a self-targeted upregulatory connection of .8 to represent replication, and an up-or downregulatory connection to a signaling protein targeted by the parasite. The parasite reproduction rate of .8 was chosen to align with previous agent based models of host parasite co-evolution (Schrom et al. 2018; Martin and Tate 2023). The immune effector (or sole effector in the case of the shared effector hosts) of the host network gains a downregulatory connection of -1 to the parasite.

#### 5. Host Life

Equation (1) is used to calculate changes in protein and parasite abundance for 150 time-steps, corresponding to the full host lifespan. For the first 50 time-steps following the equilibrium period the hosts receive developmental input, and synchronous hosts are infected. For the proceeding 100 time-steps there is no developmental input, and the asynchronous hosts are infected. We also conducted a smaller number of simulations where the two periods were reversed, so that the 100 time-steps of no developmental input occurred prior to the 50 time-steps of developmental signaling, flipping the infection timing for the asynchronous and synchronous simulations. These results are reported in the supplement and serve to validate our findings. We chose to simulate host-life using a large number of discrete timesteps to strike a balance between the continuous time biological processes take place over and computational resource limits.

#### 6. Fitness Calculation

Using data from the Host Life phase, Host fitness was calculated using the following equation:

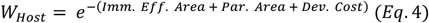

With

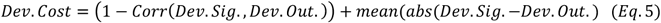

*Imm. Ef. Area*. refers to the normalized area under the curve of immune effector abundance. Immune effector area is normalized to the total length of the hosts life. *Par. Area* refers to the normalized area under the curve of parasite abundance. Parasite area is normalized to the length of time from infection start to the end of host life. Dev. Cost is our metric for capturing how closely hosts follow the developmental signal described in Equation (3). This metric captures both the mean absolute difference between host developmental effector abundance and developmental signal and the correlation between host developmental effector abundance and developmental signal. Parasite fitness was derived from the normalized area of the parasite infection, which is representative of the total number of parasites produced over the course of the infection.

#### 7. Host and Parasite Death

in each generation up to 30% of each population dies, with hosts that succumbed to infection taking priority in the death process. The 30% mark was chosen to strike a balance between overwhelming selective pressures when a greater percentage of the population died, which lead to low fitness evolutionary endpoints, and underwhelming selective pressures when a small percentage of the population died, which lead to effectively neutral evolution. The findings in this paper are potentially susceptible to large swings in this selective pressure, so results should only be directly compared to systems with approximate selection coefficients. Host deaths were first determined based on those that had failed to appropriately control their infection, meaning that *Par. Area* (defined in section 6) exceeded the death threshold of .9. This value was chosen to align with our previous work on agent-based models of host parasite co-evolution. If less than 30% of the population died due to parasite burden, surviving individuals were selected to die in a fitness-weighted manner. At random an individual was selected from the population and its chance of dying was inversely proportional to its relative fitness against the population. For example, a host that was in the 75^th^ percentile of fitness has a 25% chance of dying. This process continued until either all surviving hosts had been evaluated once, or until 30% of the population had died. In the case where greater than 30% of either population died due to poor fitness or uncontrolled infection, the excess deaths were added back to the healthy population following random selection. Population size and deaths were capped as a concession to the low fitness of initial randomly generated networks and computational expenses. Parasites deaths were determined strictly based on fitness, with the 30% of the parasite population that was least fit being selected for death in a given generation.

#### 8. Host Reproduction

survivors are picked in a fitness weighted manner to reproduce, with a host’s chance to reproduce being inversely proportional to its relative fitness against the population. A single host could produce multiple offspring in a reproductive stage. The host population was completely replenished in each reproductive stage (keeping population size constant across generations). The population in the proceeding generation was composed of all offspring all individuals that survived the death process in the current generation, along with all offspring produced in step 8. Reproduction results in the creation of a direct copy of the parent or, rarely, a mutated copy (host mutation rate: 5e-3). For hosts, a mutation could be one of the following modifications to the parent network: 1) Add a protein-protein interaction between two randomly selected proteins with a random regulatory behavior (relative probability = .25), 2) Delete a protein-protein interaction (relative probability = .25), 3) Alter regulatory coefficient (relative probability = .3), 4) Delete a protein (relative probability = .1), 5) Duplicate a protein (relative probability = .1).

#### 9. Parasite Reproduction

Parasite fitness was directly related to in host abundance, with a higher abundance indicating higher fitness. We also considered that parasites that achieve a higher within-host abundance would pass their infection on to more hosts than those that had a lower total abundance, so the number of offspring a parasite generated was dependent on their total abundance. Parasites with an abundance between 0 and .33 produced one offspring, those with an abundance between .34 and .66 produced two offspring, and an abundance greater than .66 generated 3 offspring in the next generation. Parasites reproduced until the population was completely replenished. Due to the potential for one parasite to produce multiple offspring, no parasite was guaranteed a position in the next generation in the way that surviving hosts were. Parasites reproduction directly copied the parent, or rarely a mutated copy was produced (mutation rate of 1e-2 for parasites). Parasite mutations are defined as follows: 1) randomly change the target signaling protein (relative probability = .5), 2) change the regulatory behavior on the targeted signaling protein to a new value in [-1,1] (relative probability = .5).

### Competitive simulations

Competitive simulations were used to directly evaluate the advantages and disadvantages associated with sharing an effector to accomplish multiple distinct tasks. Competitive simulations began with a burn-in period of isolated evolution, as described above, for 250, 500, or 1000 generations. During the isolated evolution phase of the simulations, populations of hosts with shared effectors (n = 250) and hosts with distinct effectors (n = 250) co-evolved with their own parasites (n = 125). When the isolated evolution period ended, the shared effector and distinct effector host populations were combined along with their respective parasite populations. Hosts competed in a fitness weighted manner for space in the next generation and competition was resolved when either one population of hosts went extinct or the competition had continued for 1000 generations.

### Resource limits on protein activation

To simulate a restriction on signaling network activity associated with limited resources, we restricted the creation of new active protein per time-step based on resource availability. When resources were scarce, we limited the generation of new active protein to .1 meaning the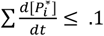 .1 for all positive changes in 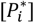. When activation of new proteins exceeded .1, all upregulatory behavior was proportionally reduced such that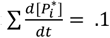 = .1. When resources were plentiful, generation of new active protein was limited to 1,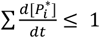. As with the scarce resource condition, when this inequality was violated by regulatory activity, all upregulatory behavior was proportionally reduced until 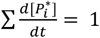.

### Host robustness

To determine if hosts with shared effectors were more susceptible to parasitic manipulation than non-pleiotropic hosts, we calculated the mean absolute difference in active immune effector abundance between intact hosts and hosts with a single signaling protein knockout. The knockout network was generated by removing a single signaling protein from the network, a standard approach when studying network robustness (Salathé and Soyer 2008; Schrom et al. 2018). Hosts were infected with a non-disrupting parasite (a parasite that could not interfere with host signaling proteins) and completed steps 2-5 described above in Simulations of Independent evolution. The mean of the absolute difference between the intact and knockout networks effector abundance was calculated and is used as a metric of the networks reliance on a specific signaling protein to produce the evolved response. See supplemental figure 1 for a diagram of the host robustness calculation process. Within a resource condition, independence of samples was determined using a Kruskal-Wallis non-parametric ANOVA and if significant differences were detected, multiple comparisons were carried out using pairwise Mann-Whitney U tests.

**Figure 1:**
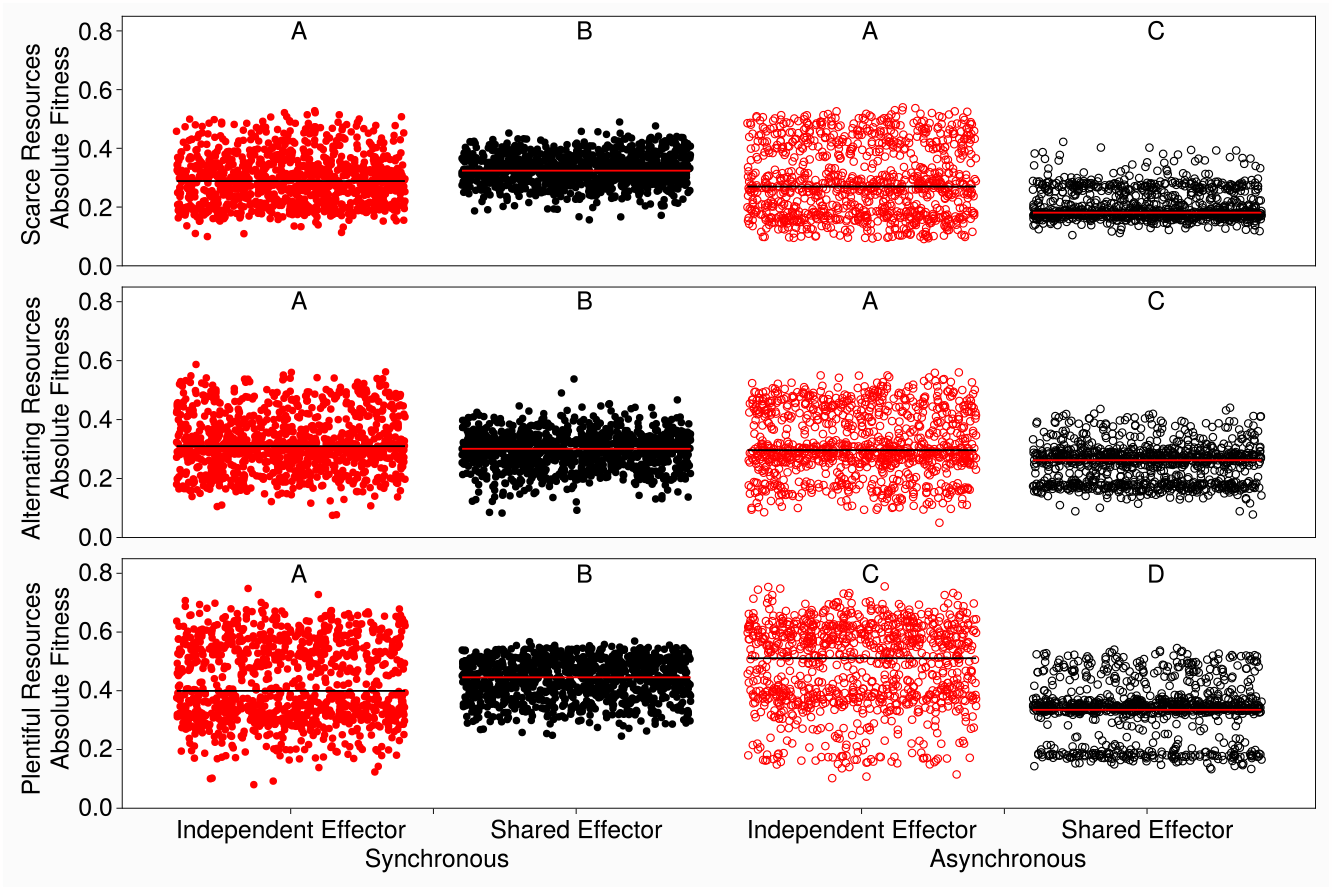
Independent effector hosts tend to be significantly more fit than shared effector hosts. Each dot represents the mean fitness of the shared effector population (black) or independent effector population (red) in the final generation of a simulation. Filled dots correspond to the synchronous signaling condition, empty dots correspond to the asynchronous signaling condition. The top row shows the scarce resource condition, the middle shows the alternating resource condition, and the bottom shows the plentiful resource condition. The y-axis shows the average absolute fitness of all hosts in the final generation of an independent evolution simulation. Within a resource condition, independence of samples was determined using a Kruskal-Wallis non-parametric ANOVA and if significant differences were detected, multiple comparisons were carried out using pairwise Signed Rank tests. Groups that share a letter are not significantly different from each other and letters are re-used between resource conditions.

### Network Features

All network features were determined for the most common host in a population at the end of independent simulations. Network connectivity was calculated by counting the number of protein-protein interactions in the immune and developmental networks and dividing that number by the number of possible connections those networks could possess. Only connections within networks were considered for network connectivity calculations and the independence of samples was determined using a Kruskal-Wallis non-parametric ANOVA. If significant differences were detected within a resource condition, multiple comparisons were carried out using pairwise Signed Rank tests. Network size was determined by counting the number of proteins present in both the immune and developmental network, the reported network size for independent effector hosts was reduced by one to adjust for the advantage of starting with 10 proteins rather than 9 in the shared effector case. The independence of samples within a resource condition were determined using a Kruskal-Wallis non-parametric ANOVA. If significant differences were detected, multiple comparisons were carried out using pairwise Signed Rank tests. Statistical comparisons of variance between samples, such as the variance in absolute fitness values attained by a population, were conducted using the variance F-test with significance threshold adjusted by Bonferroni correction.

### Quasi-Binomial Regression

To determine the probability of shared effector hosts winning a competitive simulation we calculated a binomial regression with the logistic link function using signaling timing, generations prior to competition, and resource availability as predictors of the final proportion of the population composed of shared effector hosts. Because these proportions were not strictly binomial, we adjusted the traditional regression by estimating the dispersion parameter of the quasi-binomial distribution and using this parameter to update the standard errors of the regression. The model fit was Proportion of Population that shares an effector ~ synchronous signaling + resource availability + Generations of evolution prior to competition.

## Results

### The fitness benefits of independent effectors depend on synchrony and resource availability

After the final generation of independent evolution, hosts with independent effectors attained significantly higher absolute fitness than shared effector hosts across all resource conditions when trait expression was asynchronous, as well as under variable resource conditions with synchronous signaling. Shared effector hosts attained significantly higher fitness only when immune and developmental activity were synchronous and resources were scarce (**figure 1**). Broadly, scarce resource environments producing hosts that were less fit than those in plentiful resource environments in the same signaling conditions. Hosts that evolved in the alternating resource condition attained absolute fitness levels that were between the values attained by either scarce or plentiful resource conditions. When looking at simulations where the developmental signaling period occurred in the final 50 time-steps of the simulation, the same broad trends in absolute fitness differences are observed (**figure S2**), demonstrating that these findings are robust to the order of trait expression.

We observed drastic differences in the variance around the mean of host population fitness, however. In all resource and signaling conditions, hosts with independent effectors had significantly higher absolute fitness variance as determined by the variance F-Test. The asynchronous signaling condition resulted in universally elevated absolute fitness variance, and within each signaling condition the plentiful resource condition resulted in greater fitness variance than either the alternating or scarce conditions (**table S1**).

### Shared effector hosts have a competitive advantage over independent effector hosts

Despite their lower arithmetic fitness in most scenarios, hosts with a shared effectors outcompeted hosts with independent effectors, winning more than 50% of competitive simulations in nearly all signaling and resource conditions. Shared effector hosts performed especially well when signaling was synchronous, as shown by all synchronous signaling win percentages being higher than their asynchronous counterpart. Independent effector hosts won the majority of competitive simulations when signaling was asynchronous and resources were plentiful or alternating (following 500 or 1000 generations of independent evolution). Across all resource and signaling conditions, independent effector hosts became more competitive when the burn-in period prior to competition was longer (**figure 2**).

**Figure 2:**
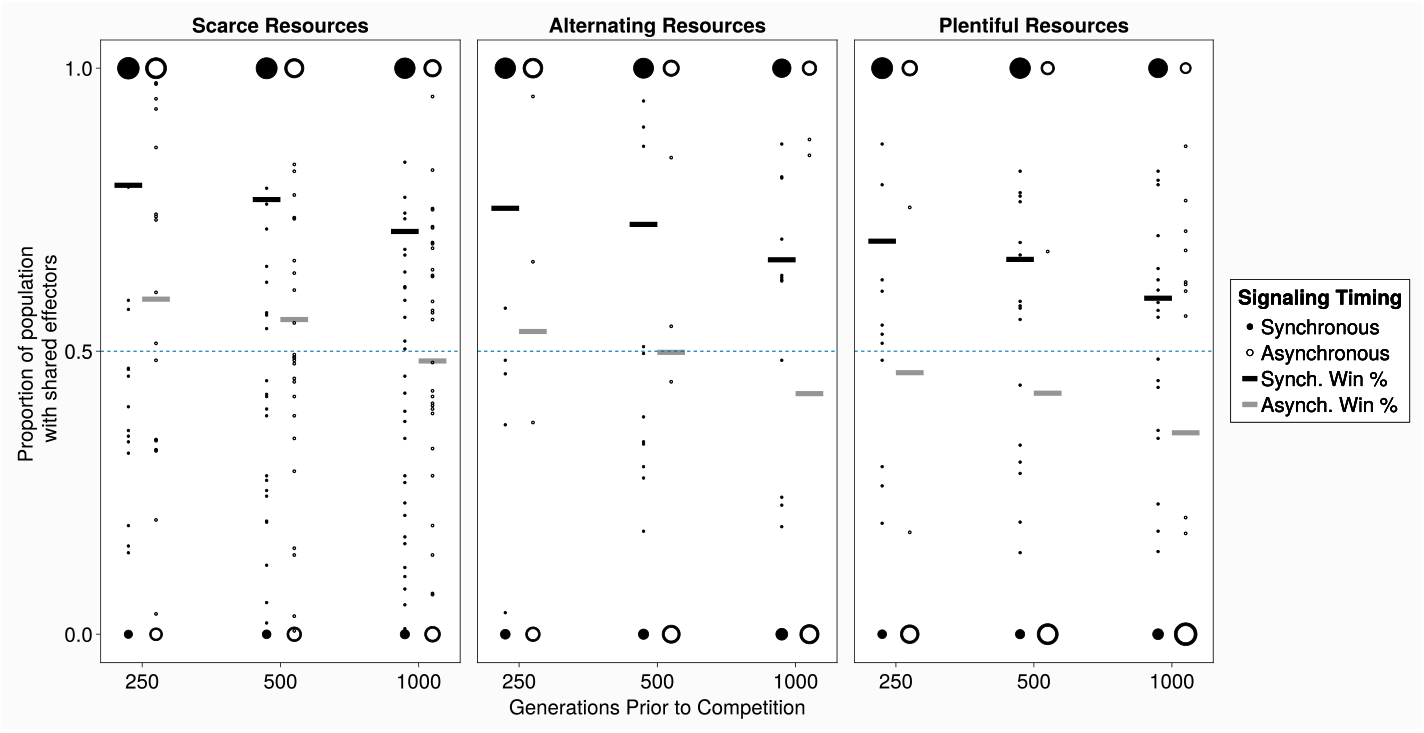
Shared effector hosts outcompete independent effector hosts in most conditions. The black lines show the predicted probability of shared effector hosts winning a competitive simulation when signaling was synchronous, the grey lines show the predicted probability of shared effector hosts winning a competition when signaling was asynchronous. In both cases, the predicted probability was determined by a quasi-binomial regression on the proportion of the population that had a shared effector in the final generation of a simulation. The y-axis shows the proportion of the host population that was composed of shared effector hosts, with the size of dots corresponding to the number of simulations that ended with that proportion. The x-axis shows the number of generations that were allotted for independent evolution prior to competition beginning.

The predicted chance of a shared effector host winning a simulation was determined using quasibinomial regression and revealed that synchronous signaling (p < 1e-99), the number of generations of evolution prior to competition (p<1e-33), and resource availability (p<1e-33) all contributed significantly to shared effector host victory. The greatest coefficient from the regression was associated with signaling timing, with shared effector hosts undergoing synchronous signaling having 2.6 times the odds of winning a competitive simulation relative to those undergoing asynchronous signaling. See supplemental table 2 for the full list of coefficients, standard errors, and odds ratios associated with the regression.

### Hosts with pleiotropic effectors are robust and highly developmentally fit

When signaling was synchronous, shared effector hosts were significantly more robust to parasite manipulation than hosts with independent effectors. When signaling was asynchronous, shared effector hosts were significantly more robust than independent effector hosts when resources were scarce, while hosts with independent effectors were significantly more robust than shared effector hosts when resources were plentiful. Universally, hosts in plentiful or alternating resource conditions were more robust than in scarce resource conditions (**figure 3**). When looking at simulations where the developmental signaling period occurred in the final 50 time-steps of the simulation, the same broad trends in host robustness are observed (**figure S3**), showing that our findings with regards to host robustness are not sensitive to the order of signaling events.

**Figure 3:**
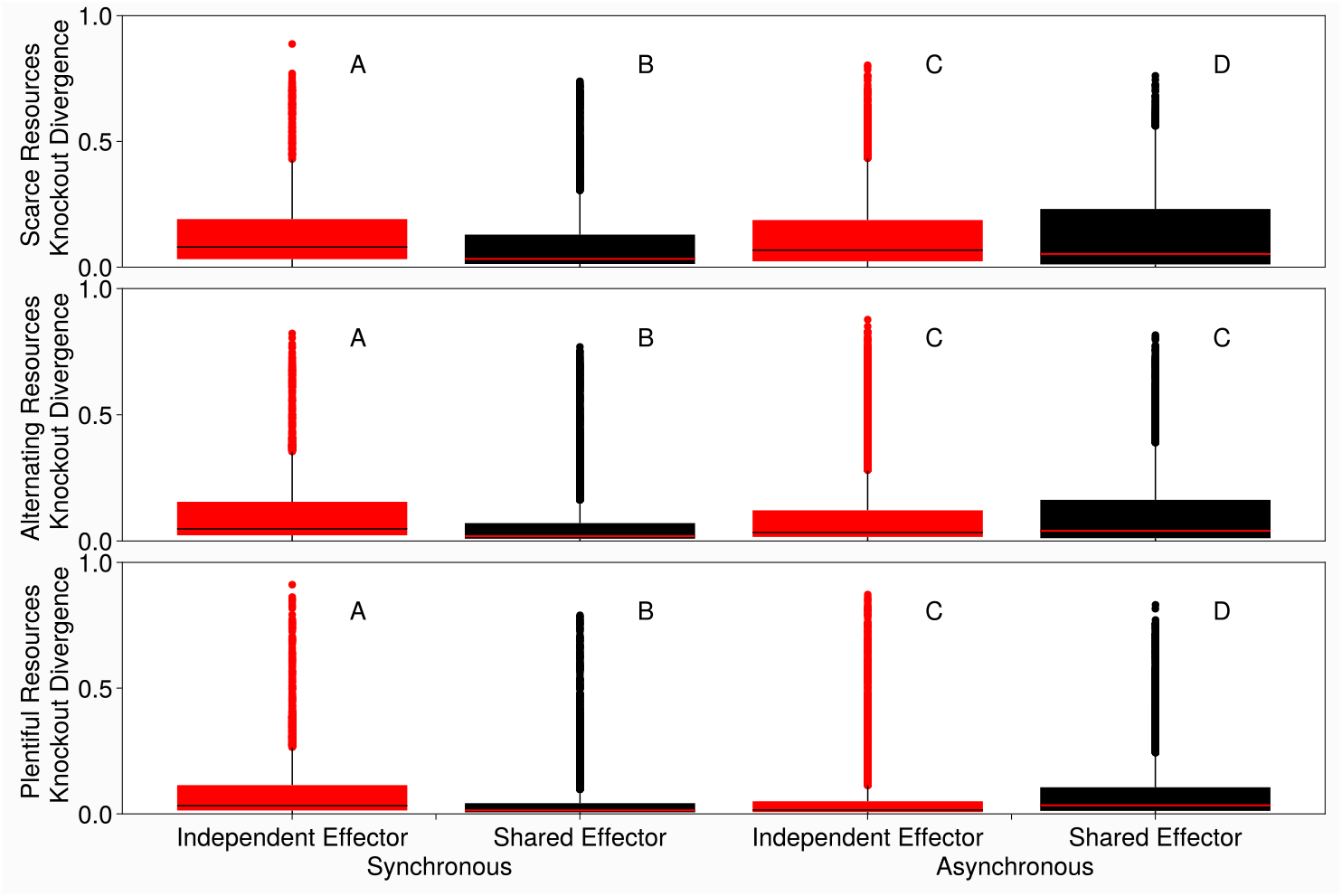
Shared effector host immune responses tend to be more robust than independent effector host immune responses. Host signaling network robustness was measured as the mean absolute difference between immune effector activity in intact and knockout hosts, using the most common hosts from the end of independent evolution simulations as the population of intact hosts. Mean absolute difference was calculated by simulating intact host infection by a non-interfering parasite and recording immune effector activity throughout the infection. A mutant host was then generated by removing a single signaling protein from the host’s immune network. The mutant then went through a simulated infection with a non-interfering parasite and the resulting immune effector activity was recorded. The absolute difference between the two immune effector abundances at each time point of the infection was then calculated and the average of all absolute differences was determined. A mutant host corresponding to a knockout of each signaling protein was created and the mean absolute difference for each was used when generating the boxplots. Higher values indicate a greater mean divergence between intact and knockout host networks, so higher scores correspond to less robust networks. The y-axis shows the mean absolute difference between the knockout immune response and the intact hosts immune response. The columns indicate the resource availability of the simulations intact hosts evolved in. Black/red lines are mean change in effector abundance following KO. Within a resource condition, independence of samples was determined using a Kruskal-Wallis non-parametric ANOVA and if significant differences were detected, multiple comparisons were carried out using pairwise Mann-Whitney U tests. Groups that share a letter are not significantly different from each other and letters are re-used between resource conditions.

Hosts with independent effectors paid lower immune costs than hosts with pleiotropic effectors across all signaling and resource conditions. These fitness cost differences were greater when signaling was asynchronous and when resources were scarce (**figure S4**). The parasite-associated fitness costs paid by shared effector hosts tended to be significantly lower than those of the independent effector hosts when signaling was asynchronous; this relationship was reversed for synchronous situations (**figure S5**). Across nearly all conditions, shared effector hosts paid significantly lower developmental costs than hosts with independent effectors, and this difference was greatest when signaling was synchronous (**figure S6**).

### Downstream pleiotropy alters evolved host network structure

While our definition of pleiotropy focused on a shared downstream module or effector (downstream pleiotropy), we allowed for the evolution of additional pleiotropic connections between the signaling proteins of the developmental and immune networks (upstream pleiotropy). In all signaling and resource conditions, hosts with independent effectors were more likely to develop upstream pleiotropic connections than shared effector hosts. This trend was significant in all conditions except when signaling was synchronous and resources were scarce (**figure 4**). The difference between the two proportions was greater when signaling was asynchronous.

**Figure 4:**
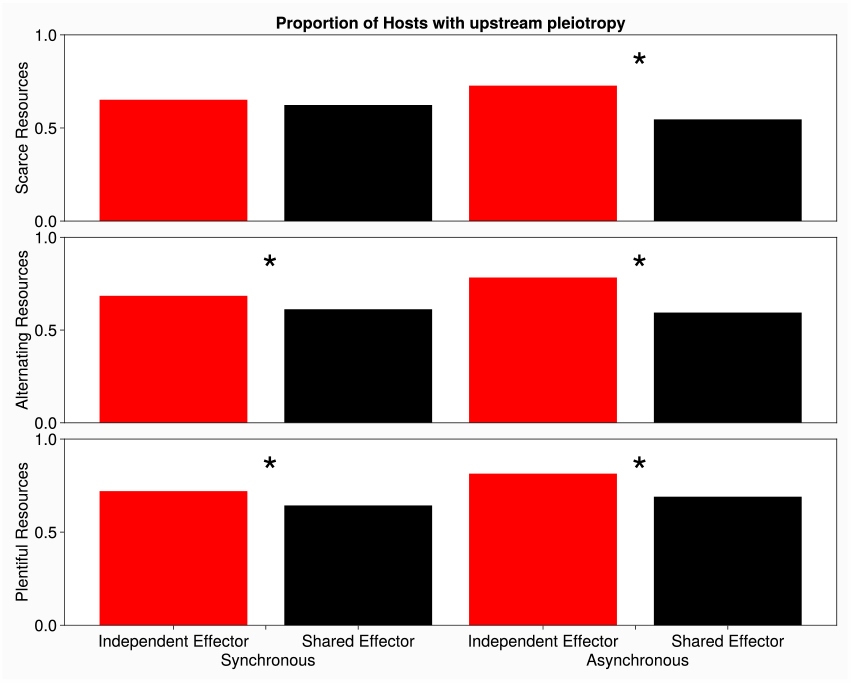
Independent effector hosts are more likely to have upstream pleiotropic connections than shared effector hosts. The y-axis shows the percent of hosts that possess at least one upstream pleiotropic connection, the x-axis groups the shared and independent effector host by the signaling condition they evolved in. The top row shows hosts that evolved in the scarce resource condition, the middle shows hosts that evolved in the alternating resource condition, and the bottom shows hosts that evolved in the plentiful resource condition. Red bars show the percent of independent effector hosts that had at least one upstream pleiotropic connection, black bars show the percent of shared effector hosts that had at least one upstream pleiotropic connection. Statistical comparisons were made between host populations within a given signaling timing and resource availability combination and significance was determined by Chi Square test, with significance threshold adjusted by Bonferroni correction. For 6 comparisons this made our alpha = .0083.

Hosts with independent effectors evolved significantly larger signaling networks than shared effector hosts across all resource conditions when signaling was asynchronous. When signaling was synchronous, shared effector and independent effector hosts tended to have similar median network sizes, but shared effector hosts had significantly greater variance in their network sizes than independent effectors hosts (**figure S7**). Hosts with independent effectors were significantly more connected than shared effector hosts across all conditions, with the difference growing more extreme when signaling was asynchronous (**figure S8**).

## Discussion

Our results suggest that pleiotropy provides a significant competitive advantage to hosts coevolving with pathogens, providing an explanation for its maintenance in signaling networks across a broad array of taxa despite the potential constraint on trait deployment and evolution (Williams et al. 2023). Our results also predict that upstream pleiotropy should be most common in hosts with no downstream pleiotropic module, highlighting a vital role for pleiotropic proteins in internetwork coordination.

We initially hypothesized that pleiotropic effectors would be detrimental to organismal fitness during synchronous signaling and that this detriment would be alleviated during asynchronous signaling because the resource allocation issue between the networks would disappear. When considering strictly the independent evolution simulations, our hypothesis is partially supported; when signaling is synchronous, shared effector hosts have lower average fitness than independent effector hosts under some resource conditions, but this finding is not universal and is even reversed when resources are scarce (**Figure 1**). This limited support of our hypothesis is further undercut when considering the results of competitive simulations, where shared effector hosts are highly successful across resource and signaling conditions (**Figure 2**). Interestingly, independent fitness is a poor predictor of competitive fitness, with shared effector hosts winning in more conditions than would be expected based strictly on the fitness values they achieve in the final generation of independent evolution (**Figures 1, 2**). Instead, signaling network robustness and the variance in absolute fitness appear to play a large role in determining which population will win a competitive simulation (**Figure 3, Table S1**). Here we have presented evidence that sharing effector modules to coordinate multiple signaling networks can be a highly fit evolutionary end point (**Figures 1, 2**) particularly when the networks are young, providing a theoretical understanding for the prevalence of downstream pleiotropy in organismal signaling networks.

As expected from well-established theory and empirical studies (Wise and Abrahamson 2005; Boots 2011; Kutzer and Armitage 2016) we found that resource availability limits absolute fitness, but both the timing of network activity and the pleiotropic nature of the effector module alter the magnitude of the penalty (**Figure 1**). The reduction in absolute host fitness due to limited resource availability seems to hinge on the relationship between resource availability and host signaling network robustness (**Figure 3**). There is significant theoretical backing for the expectation that the resource demands for a system increase as that system’s robustness increases, largely due to the costs of redundancy and the inefficiencies associated with network modularity (Kitano 2004). An empirical study from a review of the energetic economy of the human nervous system found that robust network configurations in the brain drastically increase the metabolic costs associated with connecting such robust modules (Bullmore and Sporns 2012). We found that both shared and independent effector host robustness increased when resource availability increased (**Figure 3**) and we believe that this increase in robustness can be attributed to an increase in network size as resource availability increases (**Figure S7**). This belief is supported by previous theoretical work that showed that increasing the size of the network provides a mechanism for improving network robustness (Soyer and Bonhoeffer 2006). The scarce resource condition would then prevent hosts from assuming robust, but less resource efficient, network configurations (**figures 3, S7**) and this resource derived constraint could then be reduced or avoided entirely by deploying a shared effector to accomplish multiple tasks.

Shared and independent effector hosts experience differential fitness depending on the signaling conditions they evolve in, with independent effector host fitness often increasing during asynchronous relative to synchronous signaling scenarios. Shared effector host fitness increased globally when signaling was synchronous relative to asynchronous conditions (**Figure 1**). Independent effector host fitness seems to be elevated during asynchronous signaling relative to synchronous signaling because they can deploy their effectors in a way that maximizes efficiency in both traits; having two effectors that can follow different signaling dynamics improves fitness compared to a single effector that must compromise between optimal dynamics. Shared effector host fitness seems to be elevated during synchronous relative to asynchronous signaling conditions because their single effector is capable of simultaneously accomplishing both the developmental and immune task in a resource-conserving manner; having a single effector controlled by multiple networks is beneficial when resource constraints would cause antagonistic competition between multiple effectors. Providing theoretical backing for these findings, it is expected that one of the major reasons for the maintenance of pleiotropy across evolutionary time is a single protein being capable of successfully acting in two distinct signaling pathways that are often costimulated (Cheverud 1996). We speculate that a higher resource cost associated with using effector proteins to accomplish either job would lower the shared effector host fitness gains during synchronous signaling, but not the independent effector host fitness gains. Conflict between expensive effectors could be exacerbated by low resource availability further altering our observed results. Across resource conditions and hosts, synchronous signaling lowers the variance in host fitness (**Table S1**), which may be associated with reduced population heterogeneity as there are fewer distinct ‘genotypes’ of hosts that compose a population (Perfeito et al. 2007; Lang et al. 2013). This finding suggests that the synchronous signaling condition requires relatively simple signaling network solutions, leaving less room in the fitness landscape for population heterogeneity.

While shared and independent effector hosts achieve similar functional states, both being able to successfully respond to developmental and parasitic signaling, they evolve significantly different signaling network structures to achieve these states (**Figures S7-9**). Independent effector hosts are more likely to develop upstream pleiotropic connections than shared effector hosts (**Figure 4**), suggesting that hosts with independent effectors must develop inter-network interactions to coordinate effector activity, while shared effector hosts are less reliant on such upstream pleiotropic connections for this purpose. The coordination of traits that is accomplished by both up- and downstream pleiotropic connections has a biological basis in resource management, for example; such coordination is vital for arid plant survival in resource-poor environments (Carvajal et al. 2019). Interestingly, upstream pleiotropic connections do not alter host signaling activity in the same way that downstream pleiotropic connections do (**Figures 1, 3, S7-9**), and this is likely due to the magnitude of the interaction each represents. The degree of pleiotropy, that is the number of traits a gene interacts with, positively correlates with the degree to which a gene effects host fitness (Guillaume and Otto 2012). Taken together, our data and this theoretical understanding of pleiotropy suggest that a downstream pleiotropic module has a greater degree of pleiotropy than any single upstream pleiotropic connection. Independent effector hosts also use a greater percentage of their potential internal network connections than shared effector hosts, measured as network connectivity (**Figure S8**), potentially as a supplemental mechanism of trait coordination. Theoretical studies of biological network robustness have found that highly connected networks tend to be less robust than sparsely connected networks (May 1972; Soyer and Bonhoeffer 2006), so this abundance of connections may contribute to the robustness differences observed between the shared and independent effector populations.

It is clear from our results that there is ample evolutionary space in which hosts that share an effector module can thrive, especially when competing against hosts that do not share an effector module. What is less clear however, is why this phenomenon is occurring. Prior to these experiments we predicted that independent effector hosts should be capable of at least matching, if not exceeding, the fitness of shared effector hosts. This is true during independent evolution, but not during competitive evolution simulations. We believe that the competitive advantage of shared effector hosts in this model is as follows: shared effector hosts generally have a lower variance in population fitness, especially when signaling is synchronous (**Figure 1, Table S1**), which imparts a competitive advantage against a more fit but highly variable population (Philippi and Seger 1989). Shared effector host fitness is less variable due to their robustness, as robust immune responses allow them to consistently respond to a range of parasitic manipulations. Shared effector host robustness arises because of their effector module configuration; shared effector hosts possess significantly more connections to their effector than independent effector hosts (**Figure S9**) which positively correlates with robustness (**Figure S10**). Furthermore, shared effector hosts always have lower network connectivity than independent effector hosts that evolved under the same conditions, which is associated with robustness (Soyer and Bonhoeffer 2006). A second piece of the competitive puzzle seems to be the rate at which both populations improve their fitness. We observed that shared effector hosts more rapidly improve their initial population fitness than their independent effector counterparts, especially when signaling is synchronous (**Figure S11**), as is expected of pleiotropic adaptation when initial organismal fitness is far away from a theoretical optimum, consistent with theoretical predictions (Orr 1998) and empirical study (Hämälä et al. 2020). Together, these findings paint a picture of downstream pleiotropy being advantageous, especially in early generations, which is further supported by the increased competitive success shared effector hosts experience when the populations are allowed fewer generations to evolve prior to competition beginning. The advantage associated with sharing an effector is most pronounced in synchronous signaling conditions (**Figure S2**), contradicting the notion that pleiotropy is detrimental during resource allocation trade-offs and instead providing evidence for pleiotropic effectors being evolutionarily favorable relative to independent effectors.

This work provides a theoretical framework for studying the evolutionary interactions of multiple signaling networks and is especially appropriate for understanding how they share resources and coordinate responses. In turn, these models can help to provide potential evolutionary explanations for seemingly poorly fit behaviors or phenotypes when these phenomena are inaccessible to empirical study. For example, we started this project questioning the role of phenoloxidase as a key component of both cuticle and pathogen melanization when this configuration seemingly leads to resource allocation tradeoffs. Our findings propose that in some resource and signaling environments, the current configuration is more fit than using multiple effectors, so the supposed fitness deficit is in fact the most fit implementation of the given biological network. These findings can also help to explain how novel traits rise to prominence in a population: as new traits arise they coopt existing genetic architecture (Lenski et al. 2003; Magadum et al. 2013), and this shared architecture may eventually separate into distinct traits following duplication and subfunctionalization events (Guillaume and Otto 2012; Birchler and Yang 2022). Our work has shown that the intermediate pleiotropic state, where traits share signaling components, can be well tolerated from a perspective of organismal fitness, and allows for more rapid fitness improvements than would be observed otherwise. We expect these findings to be generalizable to other traits beyond immunity and development, as the major benefit of downstream pleiotropy is an increase in signaling network robustness compared to deploying independent signaling effectors.

Signaling network robustness is proposed to be a mechanism to reduce mutational stress on signaling networks (Masel and Siegal 2009) and the pursuit of robustness has been shown to drive the evolution of signaling network complexity (Soyer and Bonhoeffer 2006).

This work could be extended in the future to investigate how the stage-structure of trait expression, e.g. in insects with complete or incomplete metamorphosis or in species that experience variable exposure to parasites in different life stages, influences the evolution of pleiotropic signaling networks. In addition, our simulations were carried out assuming that both traits contribute equally to host fitness, but imbalanced contributions may select for entirely different competitive dynamics. Finally, this work focuses on the potential benefits or costs of pleiotropy from a host perspective, but there is substantial evidence that parasites make extensive use of pleiotropic proteins, both for their own purposes (Matos et al. 2022) and as virulence factors (Kotwal and Moss 1988; Matos et al. 2022). A similar model studying parasites that deploy proteins that have unique functions between species could then be a first step in understanding the evolutionary dynamics that lead to and result from such cross-species functionality. This work provides support for the value of a network based perspective when studying trait evolution, as we were able to provide novel insights into the evolution of shared genetic architecture with implications both for the evolution and maintenance of genetic pleiotropy as well as network level resource management strategies.

## Acknowledgements

This work was supported by the National Institute of General Medical Sciences at the National Institutes of Health (grant number R35GM138007 to A.T.T.).

## List of Supplementary materials

**Table S1:** Variance of population absolute fitness values.

**Table S2:** Coefficients of the quasi-binomial regression used to determine the predicted probability of shared effector hosts winning a competition.

**Figure S1:** Diagram of host signaling network robustness calculation

**Figure S2:** Independent effector hosts tend to be significantly more fit than shared effector hosts

**Figure S3:** Shared effector host immune responses tend to be more robust than independent effector host immune responses.

**Figure S4:** Asynchronous signaling conditions lead non-pleiotropic hosts to pay lower immune costs than pleiotropic hosts.

**Figure S5:** Resource limits, signaling conditions, and pre-competition evolutionary time all alter parasite related fitness.

**Figure S6:** Pleiotropic hosts always pay lower developmental fitness costs than non-pleiotropic hosts.

**Supplemental methods:** Calculating the marginal fitness costs of shared effector modules

## References

Abdellatif, M., F. Madeo, S. Sedej, and G. Kroemer. 2023. Antagonistic pleiotropy: the example of cardiac insulin-like growth factor signaling, which is essential in youth but detrimental in age. Expert opinion on therapeutic targets.

Adamo, S. A., and N. M. Parsons. 2006. The emergency life-history stage and immunity in the cricket, Gryllus texensis. Animal behaviour 72:235–244.

Armbruster, W. S., J. Lee, and B. G. Baldwin. 2009. Macroevolutionary patterns of defense and pollination in Dalechampia vines: adaptation, exaptation, and evolutionary novelty. Proceedings of the National Academy of Sciences of the United States of America 106:18085–18090.

Bedhomme, S., J. Hillung, and S. F. Elena. 2015. Emerging viruses: why they are not jacks of all trades? Current opinion in virology 10:1–6.

Bezanson, J., A. Edelman, S. Karpinski, and V. B. Shah. 2017. Julia: A Fresh Approach to Numerical Computing. SIAM Review 59:65–98.

Birchler, J. A., and H. Yang. 2022. The multiple fates of gene duplications: Deletion, hypofunctionalization, subfunctionalization, neofunctionalization, dosage balance constraints, and neutral variation. The Plant cell 34:2466–2474.

Bisgrove, S. R., M. T. Simonich, N. M. Smith, A. Sattler, and R. W. Innes. 1994. A disease resistance gene in Arabidopsis with specificity for two different pathogen avirulence genes. The Plant cell 6:927–933.

Bleu, J., M. Massot, C. Haussy, and S. Meylan. 2012. Experimental litter size reduction reveals costs of gestation and delayed effects on offspring in a viviparous lizard. Proceedings. Biological sciences / The Royal Society 279:489–498.

Bochdanovits, Z., and G. de Jong. 2004. Antagonistic pleiotropy for life-history traits at the gene expression level. Proceedings. Biological sciences / The Royal Society 271 Suppl 3:S75–8.

Boots, M. 2011. The evolution of resistance to a parasite is determined by resources. The American naturalist 178:214–220.

Bullmore, E., and O. Sporns. 2012. The economy of brain network organization. Nature reviews. Neuroscience 13:336–349.

Carrera, J., G. Rodrigo, V. Singh, B. Kirov, and A. Jaramillo. 2011. Empirical model and in vivo characterization of the bacterial response to synthetic gene expression show that ribosome allocation limits growth rate. Biotechnology journal 6:773–783.

Carvajal, D. E., A. P. Loayza, R. S. Rios, C. A. Delpiano, and F. A. Squeo. 2019. A hyper‐arid environment shapes an inverse pattern of the fast–slow plant economics spectrum for above‐, but not below‐ground resource acquisition strategies. The Journal of ecology 107:1079–1092.

Cheverud, J. M. 1996. Developmental integration and the evolution of pleiotropy. American zoologist 36:44–50.

DiAngelo, J. R., M. L. Bland, S. Bambina, S. Cherry, and M. J. Birnbaum. 2009. The immune response attenuates growth and nutrient storage in Drosophila by reducing insulin signaling. Proceedings of the National Academy of Sciences of the United States of America 106:20853–20858.

Garland, T., Jr, C. J. Downs, and A. R. Ives. 2022. Trade-Offs (and Constraints) in Organismal Biology. Physiological and biochemical zoology: PBZ 95:82–112.

González-Santoyo, I., and A. Córdoba-Aguilar. 2012. Phenoloxidase: a key component of the insect immune system. Entomologia experimentalis et applicata 142:1–16.

Graham, A. L., E. C. Schrom 2nd, and C. J. E. Metcalf. 2022. The evolution of powerful yet perilous immune systems. Trends in immunology 43:117–131.

Guillaume, F., and S. P. Otto. 2012. Gene functional trade-offs and the evolution of pleiotropy. Genetics 192:1389–1409.

Hämälä, T., A. J. Gorton, D. A. Moeller, and P. Tiffin. 2020. Pleiotropy facilitates local adaptation to distant optima in common ragweed (Ambrosia artemisiifolia). PLoS genetics 16:e1008707.

Hughes, K. A., and J. Leips. 2017. Pleiotropy, constraint, and modularity in the evolution of life histories: insights from genomic analyses. Annals of the New York Academy of Sciences 1389:76–91.

Husak, J. F., J. C. Roy, and M. B. Lovern. 2017. Exercise training reveals trade-offs between endurance performance and immune function, but does not influence growth, in juvenile lizards. The Journal of experimental biology 220:1497–1502.

Johnston, S. E., J. Gratten, C. Berenos, J. G. Pilkington, T. H. Clutton-Brock, J. M. Pemberton, and J. Slate. 2013. Life history trade-offs at a single locus maintain sexually selected genetic variation. Nature 502:93–95.

Kinsler, G., K. Geiler-Samerotte, and D. A. Petrov. 2020. Fitness variation across subtle environmental perturbations reveals local modularity and global pleiotropy of adaptation. eLife 9.

Kitano, H. 2004. Biological robustness. Nature reviews. Genetics 5:826–837.

Kotwal, G. J., and B. Moss. 1988. Vaccinia virus encodes a secretory polypeptide structurally related to complement control proteins. Nature 335:176–178.

Kutzer, M. A. M., and S. A. O. Armitage. 2016. Maximising fitness in the face of parasites: a review of host tolerance. Zoology 119:281–289.

Lang, G. I., D. P. Rice, M. J. Hickman, E. Sodergren, G. M. Weinstock, D. Botstein, and M. M. Desai. 2013. Pervasive genetic hitchhiking and clonal interference in forty evolving yeast populations. Nature 500:571–574.

Lenski, R. E., C. Ofria, R. T. Pennock, and C. Adami. 2003. The evolutionary origin of complex features. Nature 423:139–144.

Magadum, S., U. Banerjee, P. Murugan, D. Gangapur, and R. Ravikesavan. 2013. Gene duplication as a major force in evolution. Journal of genetics 92:155–161.

Martin, R. A., and A. T. Tate. 2023. Pleiotropy promotes the evolution of inducible immune responses in a model of host-pathogen coevolution. PLoS computational biology 19:e1010445.

Masel, J., and M. L. Siegal. 2009. Robustness: mechanisms and consequences. Trends in genetics: TIG 25:395–403.

Maslennikova, S. O., L. A. Gerlinskaya, G. V. Kontsevaya, M. V. Anisimova, S. A. Nedospasov, N. A. Feofanova, M. P. Moshkin, et al. 2019. TNFα is responsible for the canonical offspring number-size trade-off. Scientific reports 9:4568.

Matos, A. L., P. Curto, and I. Simões. 2022. Moonlighting in Rickettsiales: Expanding Virulence Landscape. Tropical medicine and infectious disease 7.

May, R. M. 1972. Will a large complex system be stable? Nature 238:413–414.

McNeil, J., D. Cox-Foster, J. Slavicek, and K. Hoover. 2010. Contributions of immune responses to developmental resistance in Lymantria dispar challenged with baculovirus. Journal of insect physiology 56:1167–1177.

Moleres, J., A. Fernández-Calvet, R. L. Ehrlich, S. Martí, L. Pérez-Regidor, B. Euba, I. Rodríguez-Arce, et al. 2018. Antagonistic Pleiotropy in the Bifunctional Surface Protein FadL (OmpP1) during Adaptation of Haemophilus influenzae to Chronic Lung Infection Associated with Chronic Obstructive Pulmonary Disease. mBio 9.

Nieman, D. C., and B. K. Pedersen. 1999. Exercise and immune function. Recent developments. Sports medicine 27:73–80.

Nijhout, H. F., and D. J. Emlen. 1998. Competition among body parts in the development and evolution of insect morphology. Proceedings of the National Academy of Sciences of the United States of America 95:3685–3689.

Orr, H. A. 1998. THE POPULATION GENETICS OF ADAPTATION: THE DISTRIBUTION OF FACTORS FIXED DURING ADAPTIVE EVOLUTION. Evolution; international journal of organic evolution 52:935–949.

Ørsted, M., A. A. Hoffmann, E. Sverrisdóttir, K. L. Nielsen, and T. N. Kristensen. 2019. Genomic variation predicts adaptive evolutionary responses better than population bottleneck history. PLoS genetics 15:e1008205.

Paaby, A. B., A. O. Bergland, E. L. Behrman, and P. S. Schmidt. 2014. A highly pleiotropic amino acid polymorphism in the Drosophila insulin receptor contributes to life-history adaptation. Evolution; international journal of organic evolution 68:3395–3409.

Parker, H. H., E. G. Noonburg, and R. M. Nisbet. 2001. Models of alternative life-history strategies, population structure and potential speciation in salmonid fish stocks. The Journal of animal ecology 70:260–272.

Perfeito, L., L. Fernandes, C. Mota, and I. Gordo. 2007. Adaptive mutations in bacteria: high rate and small effects. Science 317:813–815.

Philippi, T., and J. Seger. 1989. Hedging one’s evolutionary bets, revisited. Trends in ecology & evolution 4:41–44.

Reyes, L. H., A. S. Abdelaal, and K. C. Kao. 2013. Genetic determinants for n-butanol tolerance in evolved Escherichia coli mutants: cross adaptation and antagonistic pleiotropy between n-butanol and other stressors. Applied and environmental microbiology 79:5313–5320.

Sadhukhan, A., Y. Kobayashi, S. Iuchi, and H. Koyama. 2021. Synergistic and antagonistic pleiotropy of STOP1 in stress tolerance. Trends in plant science 26:1014–1022.

Salathé, M., and O. S. Soyer. 2008. Parasites lead to evolution of robustness against gene loss in host signaling networks. Molecular systems biology 4:202.

Scarcelli, N., J. M. Cheverud, B. A. Schaal, and P. X. Kover. 2007. Antagonistic pleiotropic effects reduce the potential adaptive value of the FRIGIDA locus. Proceedings of the National Academy of Sciences of the United States of America 104:16986–16991.

Schmid-Hempel, P. 2008. Parasite immune evasion: a momentous molecular war. Trends in ecology & evolution 23:318–326.

Schrom, E. C., J. M. Prada, and A. L. Graham. 2018. Immune Signaling Networks: Sources of Robustness and Constrained Evolvability during Coevolution. Molecular biology and evolution 35:676–687.

Schwenke, R. A., B. P. Lazzaro, and M. F. Wolfner. 2016. Reproduction-Immunity Trade-Offs in Insects. Annual review of entomology 61:239–256.

Sinervo, B., and E. Svensson. 1998. Mechanistic and Selective Causes of Life History Trade-Offs and Plasticity. Oikos 83:432–442.

Sivakumaran, S., F. Agakov, E. Theodoratou, J. G. Prendergast, L. Zgaga, T. Manolio, I. Rudan, et al. 2011. Abundant pleiotropy in human complex diseases and traits. American journal of human genetics 89:607–618.

Sobecki, M., E. Krzywinska, S. Nagarajan, A. Audigé, K. Huỳnh, J. Zacharjasz, J. Debbache, et al. 2021. NK cells in hypoxic skin mediate a trade-off between wound healing and antibacterial defence. Nature communications 12:4700.

Sonnen, K. F., and C. Y. Janda. 2021. Signalling dynamics in embryonic development. Biochemical Journal 478:4045–4070.

Soyer, O. S., and S. Bonhoeffer. 2006. Evolution of complexity in signaling pathways. Proceedings of the National Academy of Sciences of the United States of America 103:16337–16342.

Soyer, O. S., T. Pfeiffer, and S. Bonhoeffer. 2006. Simulating the evolution of signal transduction pathways. Journal of theoretical biology 241:223–232.

Tian, D., M. B. Traw, J. Q. Chen, M. Kreitman, and J. Bergelson. 2003. Fitness costs of R-gene-mediated resistance in Arabidopsis thaliana. Nature 423:74–77.

Williams, A. M., T. M. Ngo, V. E. Figueroa, and A. T. Tate. 2023. The Effect of Developmental Pleiotropy on the Evolution of Insect Immune Genes. Genome biology and evolution 15.

Wise, M. J., and W. G. Abrahamson. 2005. Beyond the compensatory continuum: environmental resource levels and plant tolerance of herbivory. Oikos 109:417–428.

