## Supplementary Materials for "Pleiotropy alleviates the fitness costs associated with resource allocation trade-offs in immune signaling networks"

Martin, R. and Tate, A.T.

**Supplemental Table 1:** Variance of population absolute fitness values. Determined by calculating the variance of the absolute population fitness in the final generation of a simulation.

| | Scarce | Plentiful | Alternating |
|---|---|---|---|
| Synch. Shared | 0.0026 | 0.0046 | 0.0039 |
| Synch. Independent | 0.0070 | 0.0185 | 0.0091 |
| Asynch. Shared | 0.0028 | 0.0085 | 0.0043 |
| Asynch. Independent | 0.0141 | 0.0203 | 0.0121 |

**Supplemental Table 2:** Coefficients of the quasi-binomial regression used to determine the predicted probability of shared effector hosts winning a competition.

| | Coefficients | Standard Error | P-value | Odds Ratio |
|---|---|---|---|---|
| Intercept | .577 | .0423 | <1e-40 | .56 |
| Synchronous Signaling | .973 | .032 | <1e-99 | 2.65 |
| Pre-competition Generations | -.0006 | 5.03e-05 | <1e-30 | .999/generation |
| Resources | -.583 | .042 | <1e-41 | .56/Unit Resource |

Notes: To interpret this data, first determine if a coefficient has a significant effect on the outcome being tested against, in our case the probability of a shared effector host winning a simulation, do this by checking the p-value column and finding the coefficients that have a p-value less than a significance threshold, we used .05. Next for categorical values (Synchronous signaling) we determine the odds ratio associated with that category by exponentiating the given coefficient, telling us that a shared effector host in the synchronous signaling condition has 2.65 times the odds of winning a simulation compared to a shared effector host in the asynchronous signaling condition. For the numerical variables (Pre-competition generations, resources) we determine the odds per unit increase of the numerical value. We see that the odds of a shared effector host winning a competition decreases by .001 per generation of pre-competition evolution, and the odds of a shared effector host winning decrease by .44 for each 1 unit increase in average resource availability.

**Supplemental Figures**

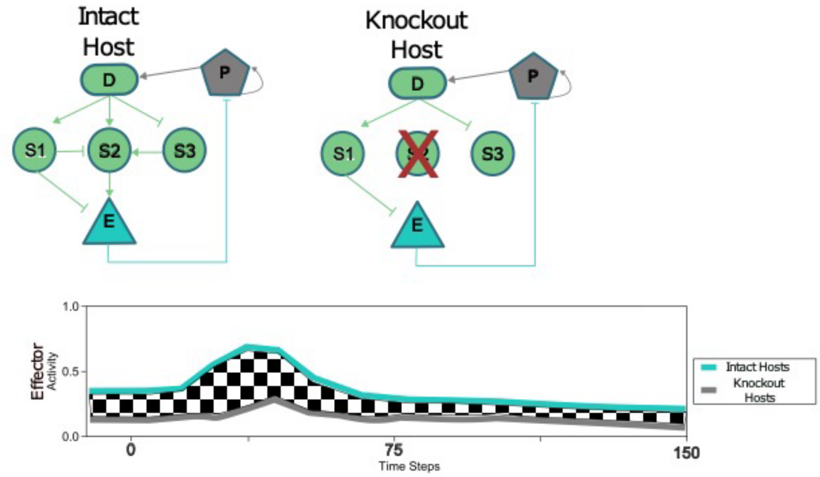


**Supplemental Figure 1:** Diagram of host signaling network robustness calculation. First intact host immune effector activity was determined during infection by a non-interfering parasite. Next, knockout immune effector activity during infection by a non-interfering parasite was calculated for a host that had a single signaling protein removed from the network (in this case signaling protein 2). The difference between these abundances (shown as the checkerboard area between the teal and grey lines) was averaged across all timesteps and recorded as the mean absolute difference between the intact and knockout host.

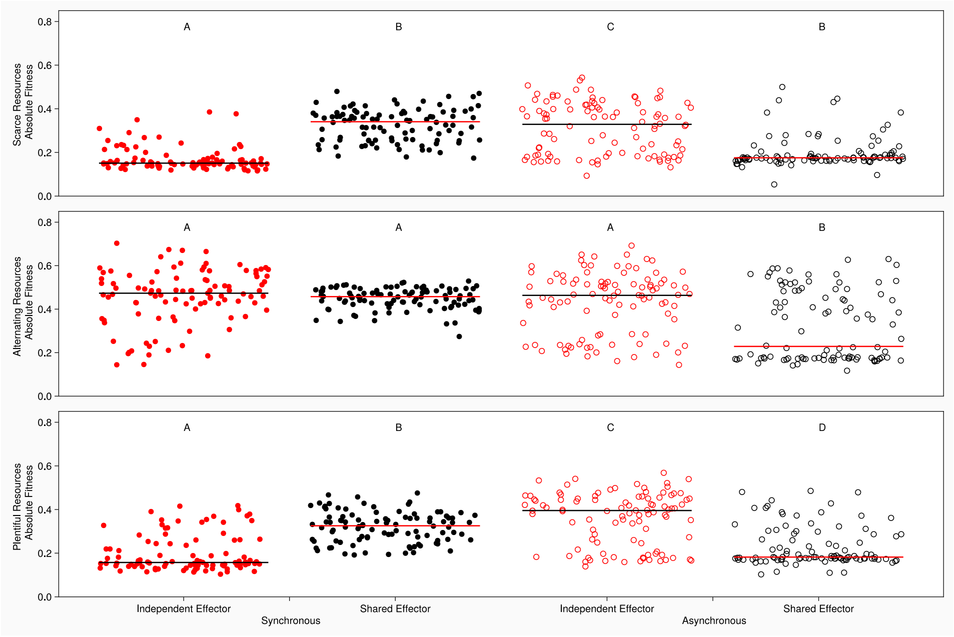


**Supplemental Figure 2:** Independent effector hosts tend to be significantly more fit than shared effector hosts. Results from 100 simulations where developmental signaling occurred after 100 time-steps of host life. Each dot represents the mean fitness of the shared effector population (black) or independent effector population (red) in the final generation of a simulation. Filled dots correspond to the synchronous signaling condition, empty dots correspond to the asynchronous signaling condition. The top row shows the scarce resource condition, the middle shows the alternating resource condition, and the bottom shows the plentiful resource condition. The y-axis shows the average absolute fitness of all hosts in the final generation of an independent evolution simulation. Within a resource condition, independence of samples was determined using a Kruskal-Wallis non-parametric ANOVA and if significant differences were detected, multiple comparisons were carried out using pairwise Signed Rank tests. Groups that share a letter are not significantly different from each other and letters are re-used between resource conditions.

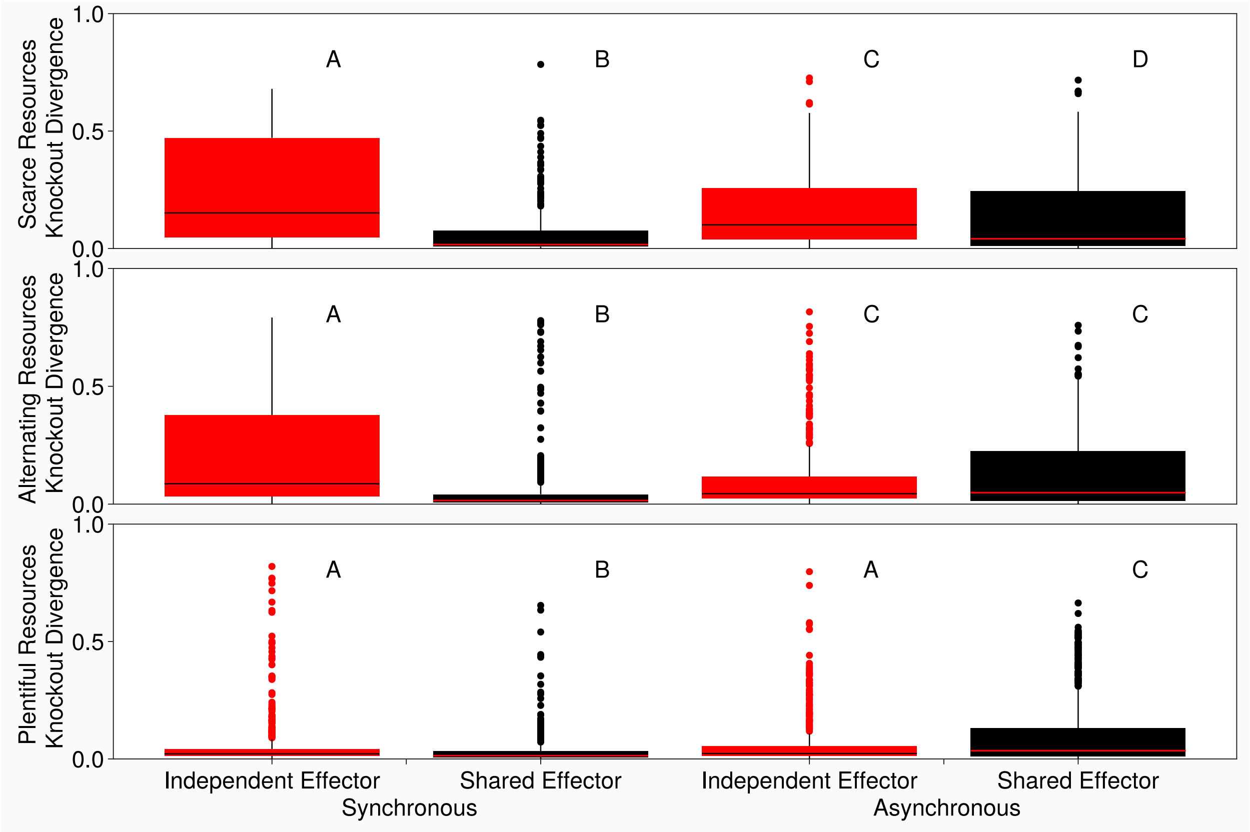


**Supplemental Figure 3**: Shared effector host immune responses tend to be more robust than independent effector host immune responses. Results from 100 simulations where developmental signaling occurred after 100 time-steps of host life. Host signaling network robustness was measured as the mean absolute difference between immune effector activity in intact and knockout hosts, using the most common hosts from the end of independent evolution simulations as the population of intact hosts. Mean absolute difference was calculated by simulating intact host infection by a non-interfering parasite and recording immune effector activity throughout the infection. A mutant host was then generated by removing a single signaling protein from the hosts immune network, the mutant then went through a simulated infection with a non-interfering parasite and the resulting immune effector activity was recorded. The absolute difference between the two immune effector abundances at each time point of the infection was then calculated and the average of all absolute differences was determined. A mutant host corresponding to a knockout of each signaling protein was created and the mean absolute difference for each was used when generating the above boxplots. Higher values indicate a greater mean divergence between intact and knockout host networks, so higher scores correspond to less robust networks. The y-axis shows the mean absolute difference between the knockout immune response and the intact hosts immune response. The columns indicate the resource availability of the simulations intact hosts evolved in. Black/red lines are mean change in effector abundance following KO. Within a resource condition, independence of samples was determined using a Kruskal-Wallis non-parametric ANOVA and if significant differences were detected, multiple comparisons were carried out using pairwise Mann-Whitney U tests. Groups that share a letter are not significantly different from each other and letters are re-used between resource conditions.

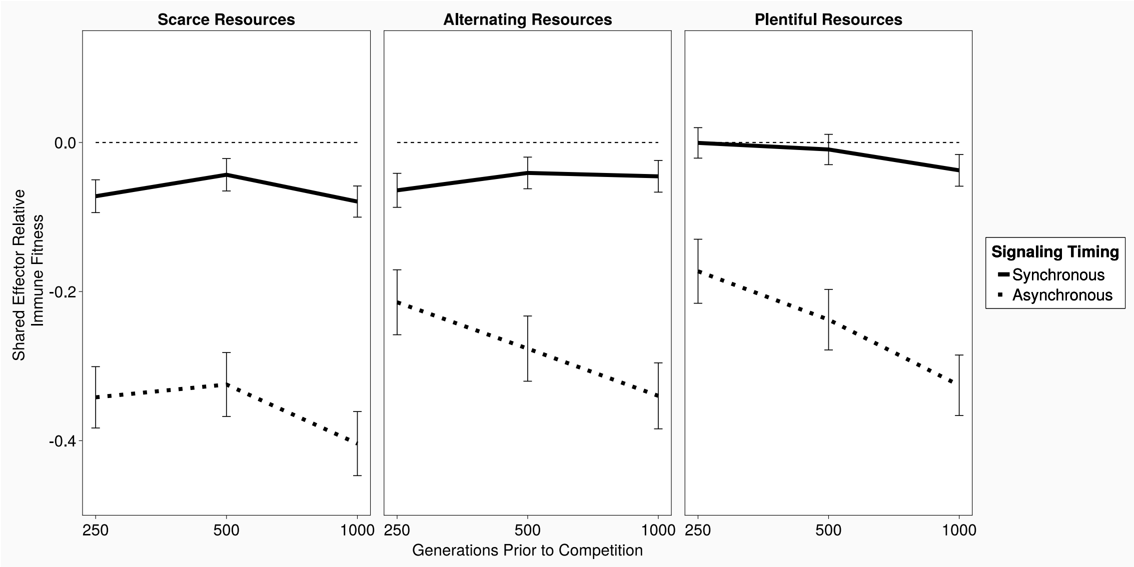


**Supplemental Figure 4:** Asynchronous signaling conditions lead non-pleiotropic hosts to pay lower immune costs than pleiotropic hosts. The y-axis is the mean difference in immune fitness costs between non-pleiotropic and pleiotropic hosts in the synchronous signaling condition (blue) and asynchronous condition (orange). The columns show the resource conditions the simulation took place in. Error bars are 95% confidence intervals around the mean, and when these bars do not include zero (dashed line) we can conclude that the true mean fitness cost difference is non-zero. When a value is below zero that indicates non-pleiotropic hosts were more fit than pleiotropic hosts, and positive values indicate that pleiotropic hosts were more fit than non-pleiotropic hosts.

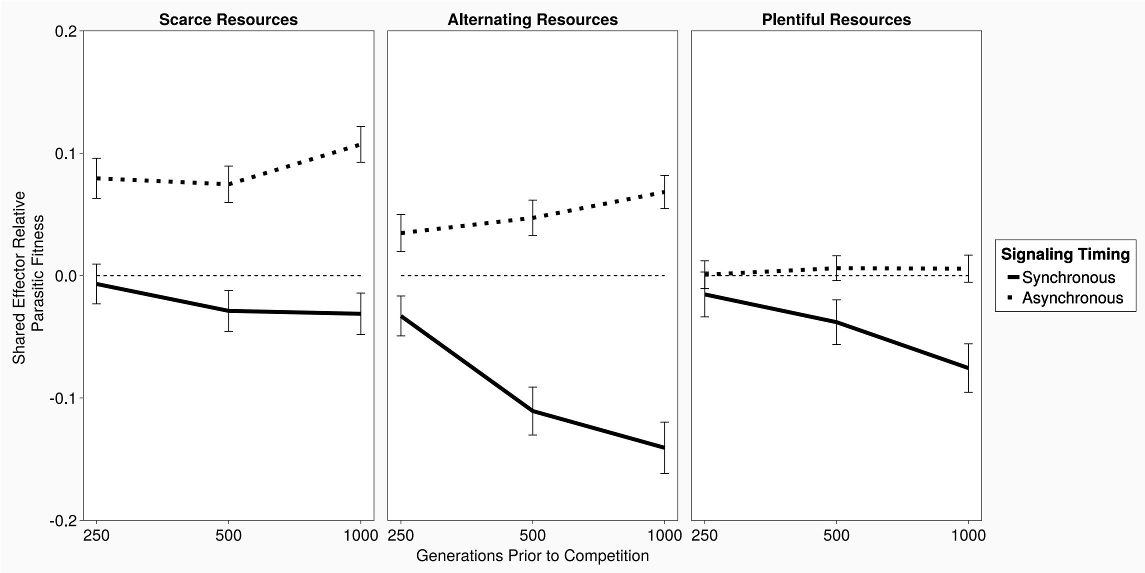


**Supplemental Figure 5:** Resource limits, signaling conditions, and pre-competition evolutionary time all alter parasite related fitness. The y-axis shows the mean difference in parasite fitness costs between non-pleiotropic and pleiotropic hosts in the synchronous signaling condition (blue) and asynchronous condition (orange). The columns correspond to the scarce, plentiful, and alternating resource conditions, from left to right. Error bars are 95% confidence intervals around the mean, and when these bars do not include zero (dashed line) we can conclude that the true mean fitness cost difference is non-zero. Values below zero indicate non-pleiotropic hosts were more fit than pleiotropic hosts, and vice versa.

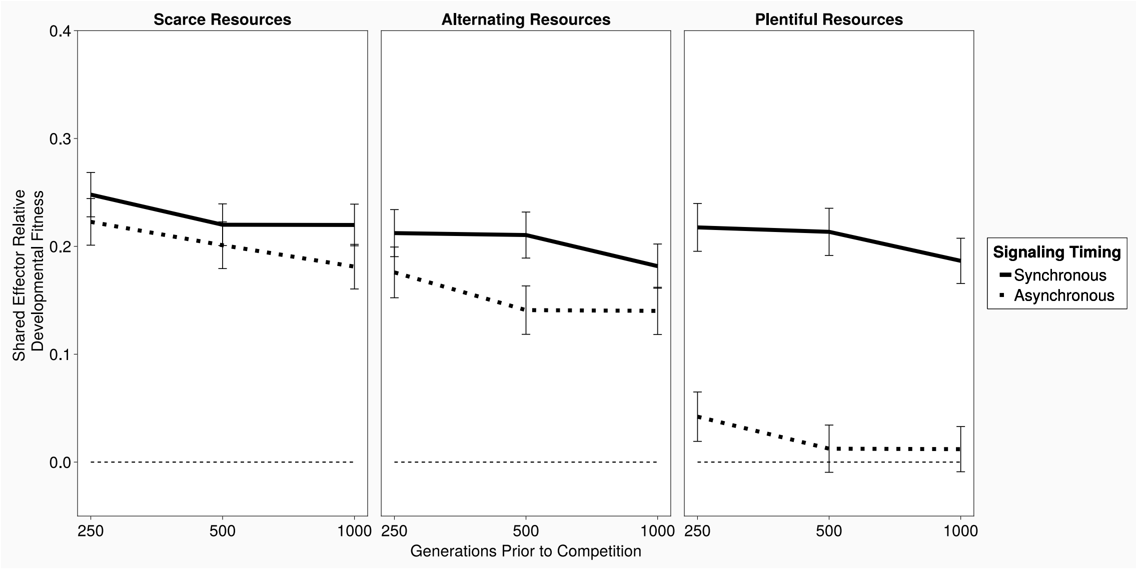


**Supplemental Figure 6:** Pleiotropic hosts always pay lower developmental fitness costs than non-pleiotropic hosts. The y-axis shows the mean difference in parasite fitness costs between non-pleiotropic and pleiotropic hosts in the synchronous signaling condition (blue) and asynchronous condition (orange). Error bars are standard error of the mean, and when these bars do not include zero (dashed line) we can conclude that the true mean fitness cost difference is non-zero. When a value is below zero that indicates non-pleiotropic hosts were more fit than pleiotropic hosts, and positive values indicate that pleiotropic hosts were more fit than non-pleiotropic hosts.

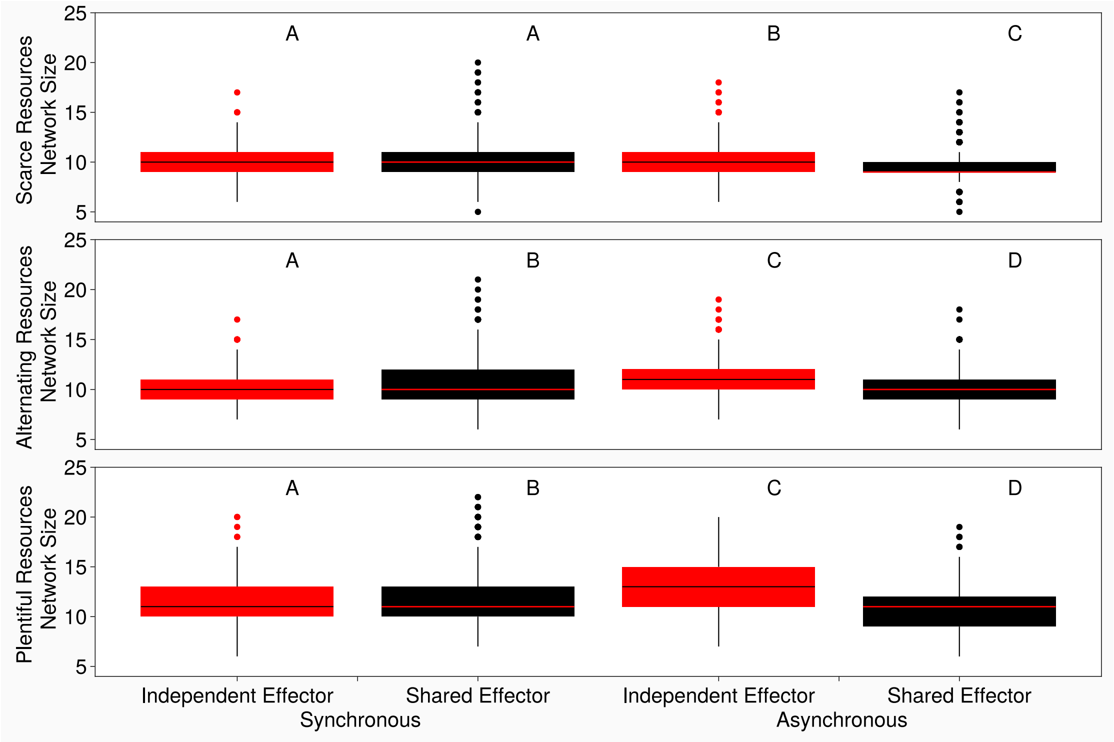


**Supplemental Figure 7:** When signaling is asynchronous independent effector host networks grow significantly larger than shared effector hosts. The y-axis shows number of proteins in the signaling network of the most common host in the last generation of a simulation. The rows show synchronous (top) and asynchronous (bottom) signaling conditions. The columns correspond to resource limits, Scarce, Plentiful, and Alternating, from left to right. Significance determined by signed rank test with significance threshold adjusted by Bonferroni correction.

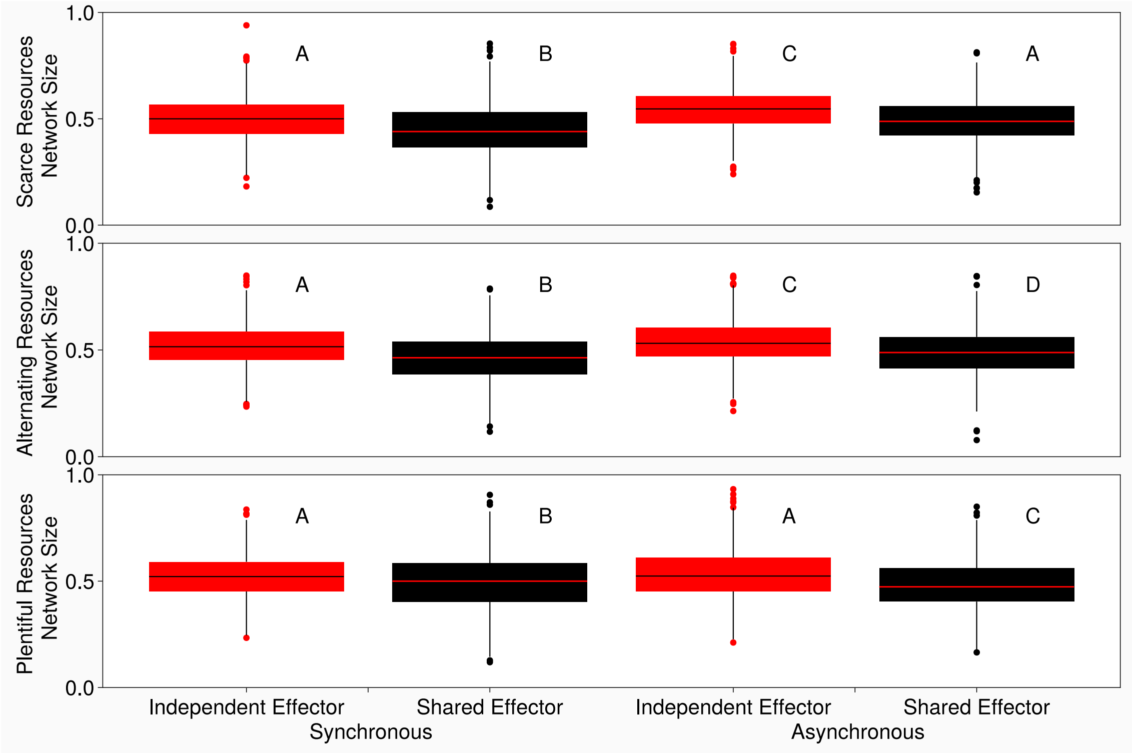


**Supplemental Figure 8:** Asynchronous signaling leads to highly connected independent effector host networks. Network connectivity, defined by $connectivity = \frac{\#\ of\ protein{-}protein\ connections}{\#\ of\ possible\ protein{-}protein\ connections}$ plots. Rows correspond to synchronous signaling (top) and asynchronous signaling (bottom) and columns correspond to the scarce, plentiful, or alternating resource conditions, from left to right. Within a resource condition, independence of samples was determined using a Kruskal-Wallis non-parametric ANOVA and if significant differences were detected, multiple comparisons were carried out using pairwise Signed Rank tests. Groups that share a letter are not significantly different from each other and letters are re-used between resource conditions.

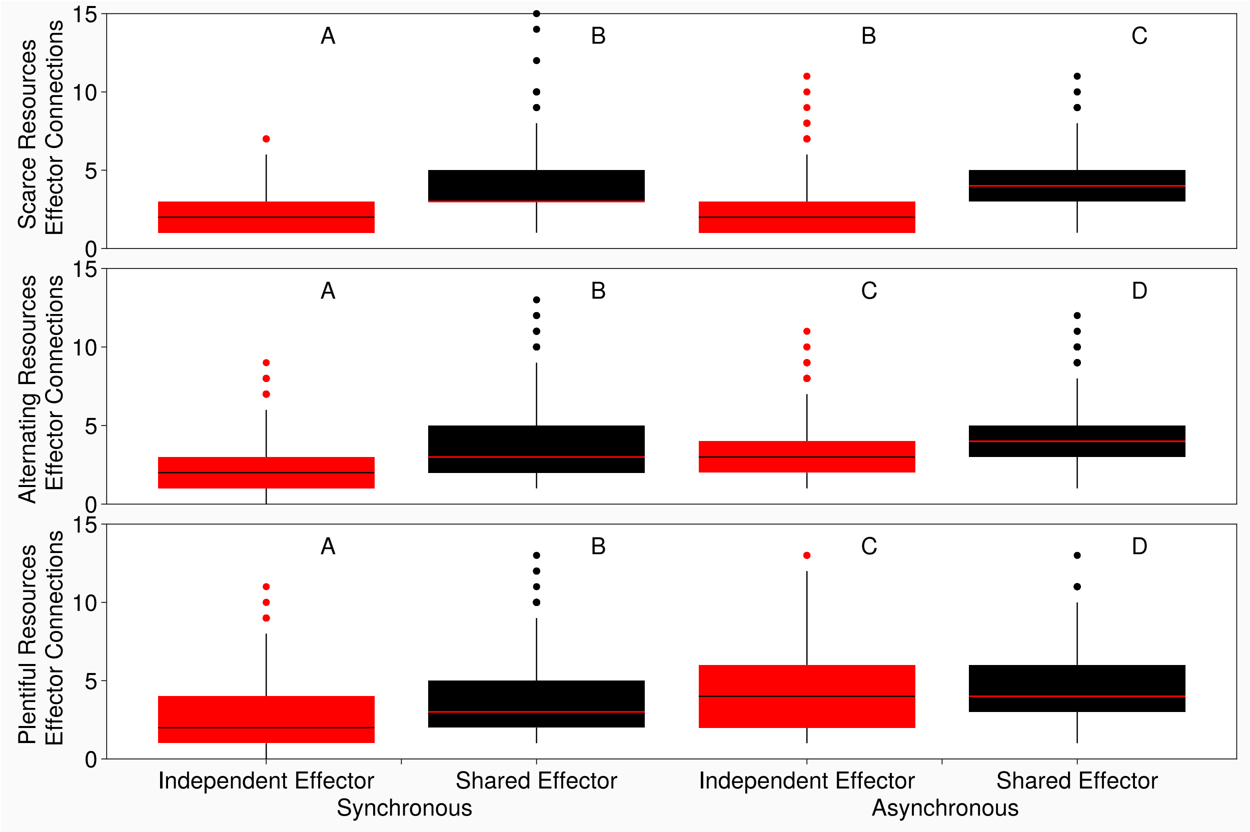


**Supplemental Figure 9:** The number of connections from signaling proteins to the immune effector (non-pleiotropic hosts) or the shared effector (pleiotropic hosts). The y-axis shows number of connections from signaling proteins to the immune or shared effector. Rows show synchronous (top) and asynchronous (bottom) signaling conditions. Columns correspond to resource limits, Scarce, Plentiful, and alternating, from left to right. Significance determined by signed rank test with significance threshold adjusted by Bonferroni correction.

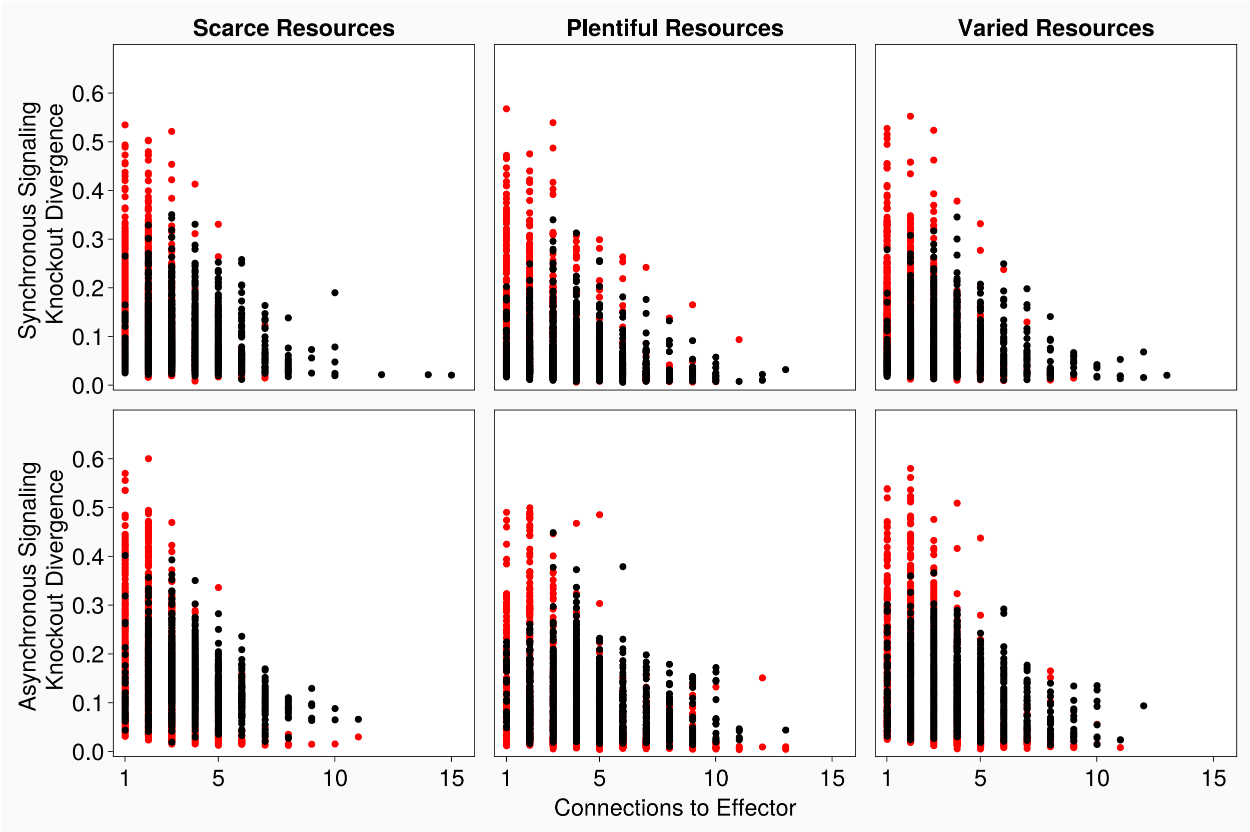


**Supplemental Figure 10:** Plotting the number of connections from signaling proteins to the immune effector versus the average divergence in effector activity due to signaling protein knockout in shared effector hosts (black) and independent effector hosts (red). The y-axis shows knockout divergence, measured as the mean absolute difference in effector level between a host with a signaling protein knockout and an intact host. The x-axis shows the number of connections from signaling proteins to the immune or shared effector. Each dot represents the average of all knockouts for a host, with hosts taken from the collection of most common hosts in a simulation.

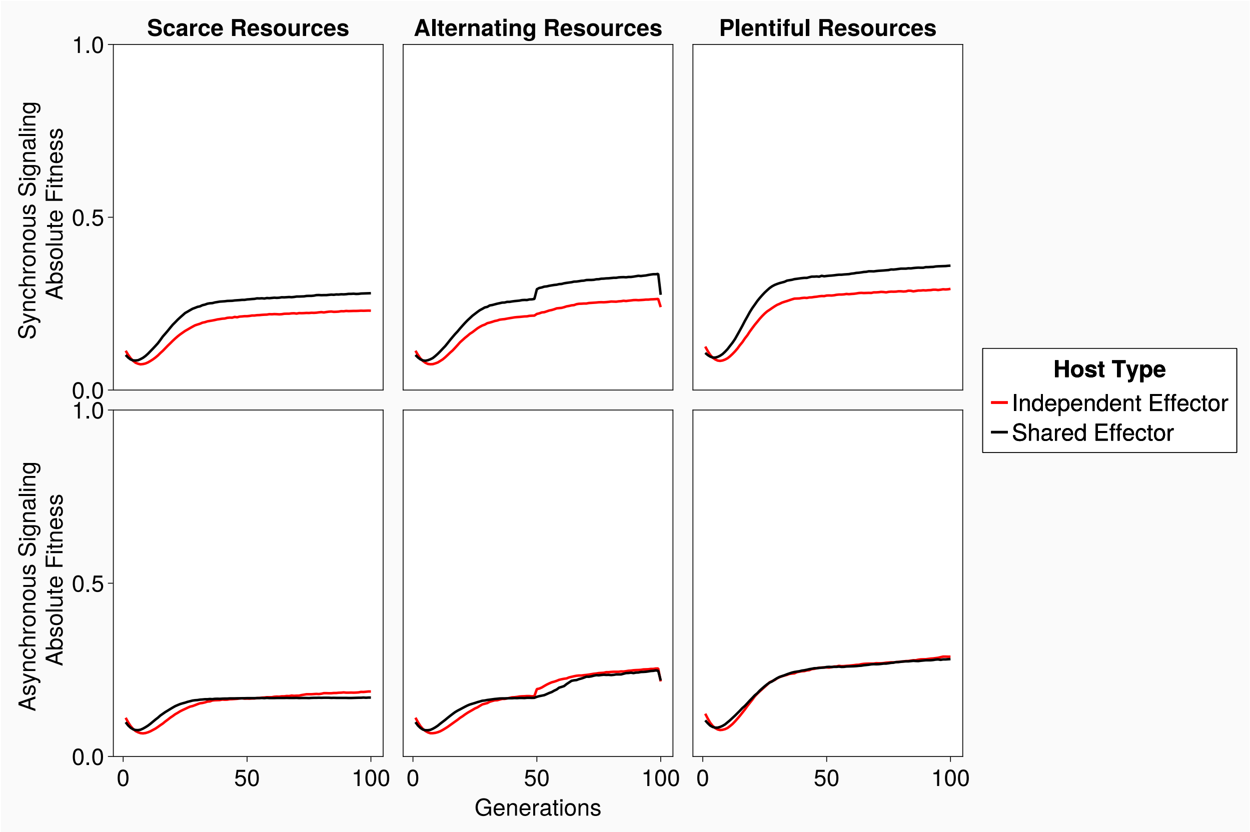


**Supplemental Figure 11:** mean population fitness through the first 100 generations of independent evolution. The top row shows fitness for hosts that evolved in the synchronous signaling condition, the bottom for hosts that evolved in the asynchronous signaling condition. The eft column shows shows that evolved in the scarce resource condition, the middle shows hosts that evolved in the alternating resource condition, and the right shows hosts that evolved in the plentiful resource condition. Black lines show shared effector host mean population fitness and red lines show independent effector host mean population fitness. The y-axis shows absolute fitness and the x-axis shows the generations of the simulation that the mean measures were taken from.

**Supplemental Methods**

### Calculating the marginal fitness costs of shared effector modules

To determine which elements of host fitness in Equation (4) most contributed to the fitness differences between Shared and Independent effector hosts we calculated the average differences between the immune, parasitic, and developmental components of fitness between the two competing populations. These differences were calculated relative to the shared effector hosts such that positive values indicated times when the shared effector hosts were more fit than the independent effector hosts and negative values indicated times when the shared effector hosts were less fit than independent effector hosts. To determine which population paid higher immune related fitness costs we would subtract the immune effector area for the shared effectors hosts from the immune effector area for independent effector hosts and take the average of these values. In an equation this would take the form

$$Immune\ Fitness\ Differences = mean(Imm.Eff.Area_{Independent\ Effector} - Imm.Eff.Area_{Shared\ Effector}),$$

substituting immune effector for parasite area or developmental costs as needed.

Here positive values indicate that on average shared effector hosts were paying a lower fitness cost for the given trait, meaning average trait specific fitness was higher than the corresponding independent effector trait specific fitness. Negative values indicate independent effector hosts were more fit. Error bars on plots correspond to 95% confidence intervals and instances where the 95% confidence intervals do not overlap the zero value indicate times when the fitness differences between the populations was significant.
